# Rational design and implementation of a chemically inducible hetero-trimerization system

**DOI:** 10.1101/2020.03.16.994277

**Authors:** Helen D. Wu, Masaki Kikuchi, Onur Dagliyan, Adam K. Aragaki, Hideki Nakamura, Nikolay V. Dokholyan, Takashi Umehara, Takanari Inoue

**Author notes:** Correspondence: Takashi Umehara for Structural Analysis, Takanari Inoue for General Correspondence.

## Abstract

Chemically inducible dimerization (CID) uses a small molecule to induce binding of two different proteins. CID tools exemplified by the FKBP/FRB/rapamycin system have been widely employed to probe molecular events inside and outside cells. While various CID tools are available, chemically inducible trimerization (CIT) has not been developed, due to inherent challenges in designing or identifying a chemical that simultaneously binds three proteins with high affinity and target specificity. Nevertheless, by introducing a third recruitable component, CIT could enable versatile applications. Here, we devised the CIT by rationally splitting FRB and FKBP. Based on cellular and structural datasets, select split pairs of FRB or FKBP underwent efficient trimerization with full length FKBP or FRB, respectively, upon addition of rapamycin. We also demonstrated its potential for cellular applications by rapidly inducing tri-organellar plasma membrane-ER-mitochondria junctions, and by perturbing intended membrane lipids exclusively at the plasma membrane-ER membrane contact sites. By conferring one additional condition to what is achievable with CID, CIT expands the types of manipulation in single live cells, to address cell biology questions otherwise intractable, and engineer cell functions for future synthetic biology applications.

## Introduction

Chemically inducible dimerization (CID) uses a bifunctional small molecule to bring together two proteins with high affinity, specificity and fast kinetics^1,2^. CID systems such as the rapamycin-dependent FKBP-FRB heterodimerization (**Fig. 1a**) can exert tight spatiotemporal control over local protein concentrations by oligomerization^3^ or by rapidly translocating a protein of interest to or from a desired site of action^4, 5^; translocation-based and other CID-based strategies have enabled signal parsing of a given protein from the noise of other cell signaling pathways, leading to discoveries in cell migration, endocytosis, transcription, and translation^1, 2, 6^. The seconds-to-minutes timescale that CID operates at offers a further advantage – CID strategies are less susceptible to genetic compensation that sometimes plague knockdown (KD) or knockout (KO) methods. When used as a building block for rewiring circuits in a bottom-up approach, CID systems have created fast, translation-independent synthetic logic gates in mammalian cells^7^. Over the years, CID has become a widely-used, easily generalizable chemical biology tool with promising clinical impact^8^.

**Figure 1:**
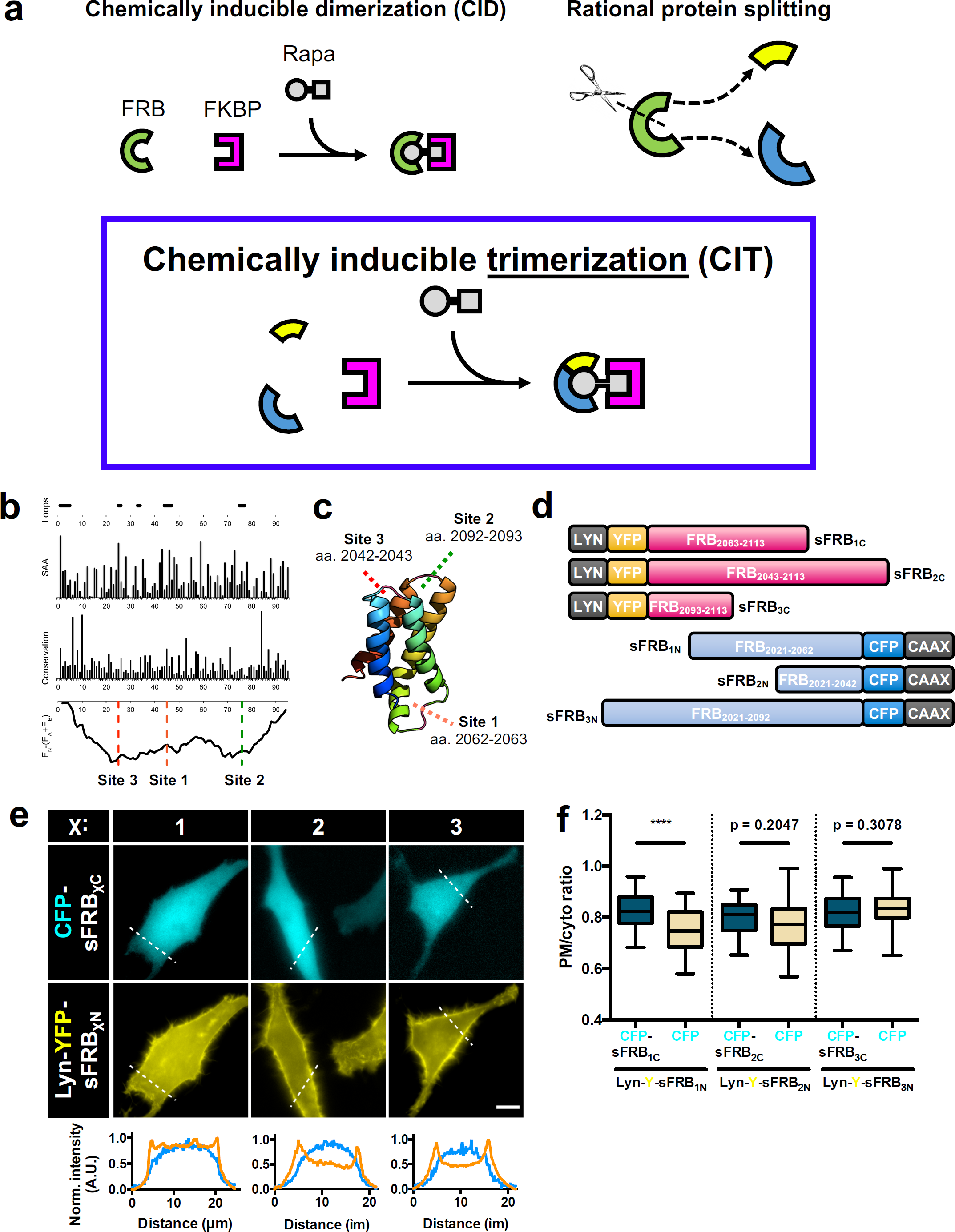
Rational CIT design and sFRB overexpression in cells. (a) CID proteins FRB (green, arc) sites generates FRB protein halves (yellow and blue). Split FRB halves and FKBP form three components of CIT. Rapamycin brings them together. (b) Computationally analysis to determine split sites for FRB domain only. Loop areas with high solvent accessibility area (SAA), low conservation and favorable split energy profiles were chosen. (c) FRB structure (PDB ID: 1AUE) consists of 4 α-helices; split sites 1, 2 and 3 all reside in linker regions between α-helices. (d) Examples of sFRB fusion protein design. Lyn or CAAX motifs target sFRBs to the PM. (e) Co-overexpression of sFRB protein pairs in HeLa cells. N-terminal sFRB halves (sFRB_1N_, sFRB_2N_, sFRB_3N_) are PM localized by Lyn and C-terminal sFRB halves (sFRB_1C_, sFRB_2C_, sFRB_3C_) are cytosolic. Linescans below show normalized PM (orange) and cytosolic (blue) signal intensity. (f) Pre-association between sFRB halves. PM-to-cytosolic ratios derived from linescans of cytosolic sFRB_1C_, sFRB_2C_, and sFRB_3C_ co-expressed with PM-targeted sFRB_1N_, sFRB_2N_, and sFRB_3N_ respectively compared to CFP alone (from left to right: *n* = 39, 33, 27, 39, 36, and 26 cells; 3 experiments). Scale bar, 10 μm.

While there have been hundreds of applications of CID, no inducible trimerization system has been developed to our knowledge. Examples of transient or stable trimerization in biology include trimeric G proteins^9^, inner ear epithelial cell junctions^10^, and MHC-antigen-TCR complexes governing T-cell selection^11^. Engineering chemically inducible trimerization (CIT) could address biologically relevant questions, and such a system would expand in general the palette of what is achievable with chemical biology.

In theory, the simplest CIT system should comprise of three unique protein components that can be brought together by a tri-functional small molecule. However, trimerization would be difficult to achieve, as recruiting an additional protein doubles the possible bound states of the small molecule compared to dimerization (**Supplementary Fig. 1**). Efficient trimerization requires that the trimeric complex be the most favorable state over other less bound configurations. For CID, the secret to its high efficiency over a large range of rapamycin concentrations is positive cooperativity: When one CID protein binds rapamycin, the resulting rapamycin-protein complex binds the second protein with thousand-fold higher affinity than rapamycin alone^12^. This unique cooperativity is rapamycin-dependent, and mediated through electrostatic interactions at the FKBP-rapamcycin-FRB interface^13^. Because engineering *de novo* proteins with appropriate small molecule-dependent surface interactions would be highly challenging, we instead exploited the special properties of the FKBP-FRB CID system to achieve trimerization.

We report the generation of a novel CIT system whose components comprise split FRB and FKBP proteins **(Fig. 1a)**. Determined with the aid of computational analysis, candidate split proteins were characterized for their efficacy of trimerization in cells. Formation of the trimer *in vitro* was assessed by X-ray crystallography. CIT was used to target cytosolic proteins to regions of close inter-organelle membrane junctions and to induce tri-organellar ER-plasma membrane-mitochondria contacts. Finally, we used CIT to locally alter phosphoinositide composition of ER-plasma membrane junctions to affect local lipid homeostasis. Altogether, CIT has small components that trimerize with fast kinetics in a rapamycin-inducible manner, expanding the variety of perturbations possible in the chemical biology toolkit.

## Results

### Design of split FRB and FKBP

We narrowed down candidate split site regions of FRB and FKBP by avoiding regions forming secondary structures, evolutionarily-conserved amino acids or residues crucial for CID functionality. We used FRB (residues 2021–2113 of human mTOR) with the T2098L mutation, which makes FRB otherwise unstable unless bound to chemical ligands or FKBP^14^, an advantageous trait for low recombination potential after protein splitting (**Fig. 1b**). In splitting FRB, we avoided its four alpha helices, crucial residues for FRB-FKBP-rapamycin stabilization (L2031, S2035, Y2038, F2039, T2098L, W2101, D2102, and Y2105)^13^, and regions of high conservation in the 3 FRB loops (**Fig. 1c** and **Supplementary Fig. 2a,b**). We then applied the SPELL algorithm^15^ to pinpoint candidate split sites in FRB through computational analysis. All the loops selected for splitting (**Fig. 1b,c** and **Supplementary Fig. 2a**) show low evolutionary conservation with Kullback-Leibler conservation scores less than 2 and have solvent accessible area values higher than 30 Å^2^ (**Fig. 1b**)^15,16,17^. We also generated the split energy profile by computing the energy of the split parts relative to the native energy of the intact protein (**Fig. 1b**); the three loops reside in two major energy wells, which suggest splitting at these sites would generate split proteins that are less likely to pre-associate when co-expressed in the absence of rapamycin ^15^. Split sites for FKBP were also determined in this manner (**Supplementary Fig. 4a,b**). Hereafter, split FRB or FKBP proteins are abbreviated as sFRB or sFKBP. Each split site is numbered in terms of likelihood of success based on the computational analysis; sFRB_1_, sFRB_2_ and sFRB_3_ refer to pairs of sFRB proteins split at sites 1, 2 or 3, with split site 1 most likely to succeed. The N- or C-terminal half of a given sFRB protein pair is sFRBχ_N_ or sFRBχ_C_, χ being the split site number. A similar naming convention was used for split FKBP.

To determine whether split proteins can be stably expressed, we generated plasmids of N-terminal fluorophore-tagged sFRB proteins and observed cytosolic expression in HeLa cells (**Supplementary Fig. 2c**). In some cases, split proteins show ER accumulation, which may be attributed to instability in protein folding. We also generated a series of Lyn or CAAX membrane-tagged sFRB fusion proteins, all of which localized to the plasma membrane (**Fig. 1d**). To determine whether there is appreciable pre-association between sFRB pairs, we used the Lyn motif to target sFRBχ_N_ to the plasma membrane (PM), observed the localization of corresponding cytosolic sFRBχ_C_ across all three split sites (χ = 1, 2, 3), and determined PM/cytosol signal ratio of the cytosolic protein (**Fig. 1e,f**). Higher PM/cytosol ratios occurred between Lyn-Y-sFRB_1N_ and CFP-sFRB_1C_ compared to Lyn-Y-sFRB_1N_ and CFP alone (0.83 ± 0.01 vs. 0.75 ± 0.01, p < 0.0001). In contrast, cytosolic CFP-sFRB_2N_ and sFRB_3N_ PM/cytosol ratios did not differ from CFP alone (0.80 ± 0.01 vs. 0.77 ± 0.01, 0.82 ± 0.01 vs. 0.84 ± 0.01, respectively) with the same experimental setup, suggesting the sFRB_1_ pair has higher tendency to pre-associate. This approach to designing split sites, validating cytosolic protein expression, and generating membrane-anchored protein was also applied to sFKBP.

### Characterization of split FRB

The sFRB protein pairs were tested for their ability to bind FKBP in a rapamycin-inducible manner to achieve CIT. We co-expressed cytosolic FKBP with PM-localized sFRB pairs in HeLa cells. If functional, rapamycin-induced FKBP binding to sFRB pairs would shift FKBP localization from the cytosol to the PM (**Fig. 2a**).sFRB pairs were localized on the PM with Lyn and CAAX tags (**Fig. 1d**). Upon rapamycin addition, all three sFRB pairs indeed recruited cytosolic mCh-FKBP to the PM, comparable to FKBP recruitment by full length (FL) FRB (**Fig. 2b**). Changes in cell linescans reflect the cytosol-to-PM localization shift pre- and 9 mins post-rapamycin administration (**Fig. 2b**). Percent change in mCh-FKBP PM/cytosol ratios derived from linescans was positive for all sFRB pairs tested; values were not significantly different between sFRB_1_, sFRB_2_, and sFRB_3_ versus full length FRB (**p = 0.613, 0.057, 0.579 respectively, Fig. 2c**). However, no sFRB_C_ half could recruit FKBP to the PM without co-expression of its cognate sFRB_N_ half and vice versa (p < 0.0001 for all cases, **Fig. 2c**). These results suggest that PM-localized sFRB halves are as efficient as full length FRB in recruiting FKBP in the presence of rapamycin, but each sFRB half cannot independently recruit FKBP. However, in the absence of FKBP overexpression, sFRB halves can still dimerize (**Supplementary Fig. 3**), as shown by rapamycin-dependent recruitment of cytosolic sFRBχ_C_ to PM-localized Lyn-YFP-sFRBχ_N_ (χ = 1, 2, 3). This is most likely due to endogenous FKBP-rapamycin complexes in the cell ^18^, and/or due to rapamycin alone bringing sFRB pairs together. sFKBP pairs were also screened, and 3 out of 4 split sites (χ = 1, 2, 3) produced functional sFKBP CIT systems (**Fig. 3f and Supplementary Fig. 4**).

**Figure 2:**
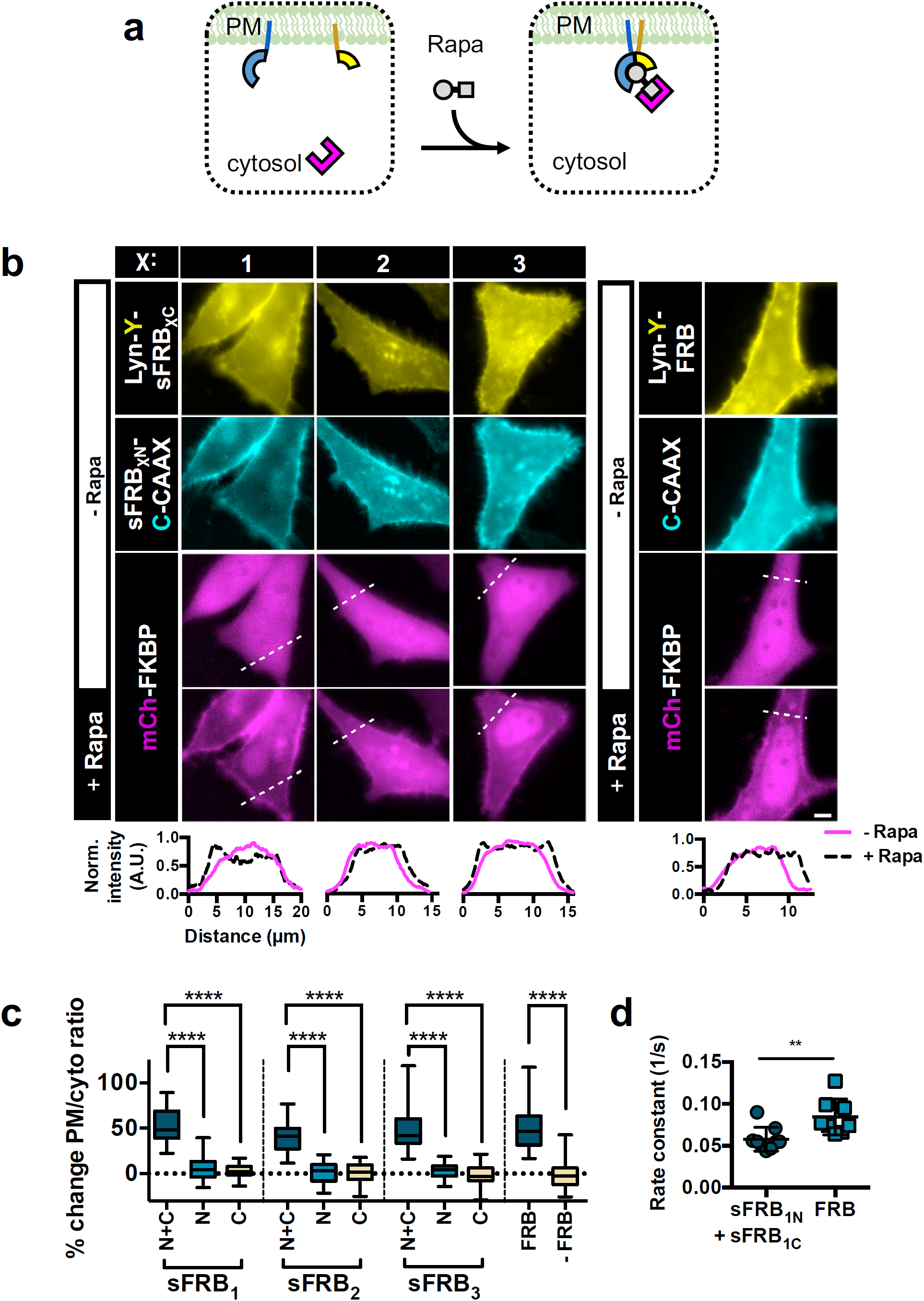
Characterization of FKBP recruitment extent and kinetics by sFRB after rapamycin addition. (a) Experimental setup for FKBP recruitment efficacy by sFRBs. sFRB N-terminal and C-terminal halves are PM-targeted by CAAX or Lyn respectively and co-expressed with FKBP. Cytosol-to-PM translocation by FKBP indicates successful recruitment. (b) Translocation of FKBP from cytosol to PM in HeLa cells co-expressing PM-targeted sFRB_1_, sFRB_2_, sFRB_3_ pairs and full length FRB. Pre-rapamycin; top 3 rows. 9 min post-rapamycin; bottom row. Linescans show corresponding normalized intensity of mCh-FKBP signal. (c) Percent change in mCh-FKBP PM-to-cytosolic ratio pre- and post-rapamycin addition. FKBP recruitment by sFRB pairs (N + C) compared to N or C-terminal halves only (N or C), full length FRB or no FRB in FKBP recruitment (from left to right: *n* = 31, 29, 24, 34, 20, 24, 27, 23, 34, 47, and 43 cells; 3-6 experiments). (d) FKBP recruitment rate constants derived from exponential regression of cytosolic FKBP signal decay imaged over 10 s intervals. sFRB_1C_ + sFRB_1N_ and FRB groups use identical plasmid constructs as leftmost and rightmost columns respectively in (b) (from left to right: *n* = 9 and 8; 4-5 experiments). Scale bar, 5 μm.

**Figure 3:**
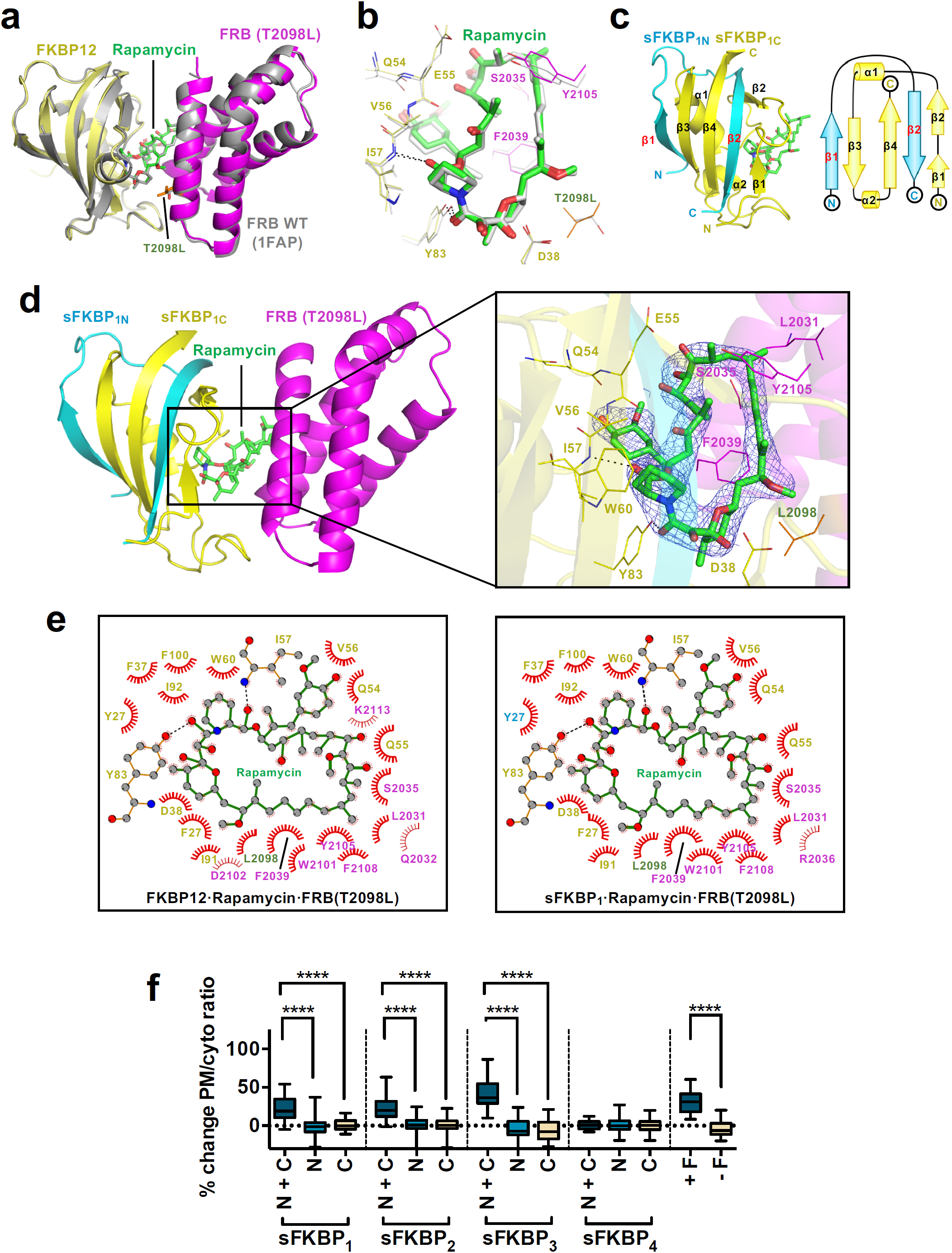
Crystal structure of the split and unsplit FKBP·rapamycin·FRB (T2098L) complexes. (a) Superimposition of FKBP12·rapamycin·FRB (T2098L) complex crystal structure (yellow, green and magenta) with that of FKBP12·rapamycin·FRB (wild type; PDB ID: 1FAP, gray). T2098L is shown in orange. (b) Zoom-in view of the rapamycin-interacting region of FKBP12·rapamycin·FRB complexes (color-coded as in panel a). Hydrogen bonds are shown as dashes. (c) Ribbon (left) and topology (right) diagrams of the sFKBP_1N_ (cyan), FKBP_1C_ (yellow) and rapamycin (green) complex. (d) Crystal structure of sFKBP1·rapamycin·FRB (T2098L). Left, sFKBP1 (cyan and yellow), rapamycin (green) and FRB (T2098L, magenta) is depicted as a ribbon diagram. Right, Electron density of rapamycin. Rapamycin is depicted as a stick model with a 2Fo-Fc electron density (blue mesh) contoured at 1.5 σ. Residues of sFKBP_1C_ (yellow) and FRB (T2098L) (magenta) involved in the interaction with rapamycin are labeled and shown as a line model. (e) LIGPLOT diagrams showing critical contacts around rapamycin. Left, FKBP12·rapamycin·FRB (T2098L). Right, sFKBP_1_·rapamycin·FRB (T2098L). Rapamycin (green) and key residues of unsplit (left) and split (right) FKBP proteins (yellow; Y27 of sFKBP_1N_ in cyan) and those of FRB (magenta; T2098L in orange) are depicted in a ball and stick model. Carbon, nitrogen and oxygen are gray, blue and red balls, respectively. Hydrogen bond and hydrophobic interaction is shown in a black dashed line and a red arc of a circle, respectively. (f) Percent change in FRB-mCh PM-to-cytosolic ratio pre- and post-rapamycin addition. Each sFKBP pair (N+C) is compared with N or C-terminal halves only (N or C), full length FKBP or no FKBP in FRB recruitment (from left to right: *n* = 21, 31, 38, 32, 39, 38, 28, 27, 21, 28, 33, 32, 29, and 31 cells; 3 – 13 experiments).

The kinetics of cytosolic FKBP recruitment to the PM was determined from exponential regression of cytosolic FKBP intensity decay during translocation. Cytosolic FKBP signal decay was captured by confocal imaging acquisition at 10 s intervals. The rate constant of cytosolic FKBP signal decay with the full-length FRB was significantly higher than with the sFRB_1_ pair (0.085 ± 0.0075 s-1 versus 0.058 ± 0.0047 s-1, p = 0.008; **Fig. 2d**), indicating that sFRB_1_ has slower FKBP recruitment kinetics upon rapamycin addition, although the timescale of FKBP recruitment by the sFRB_1_ pair still falls within the minutes range. CID is practically irreversible in cells due to low *koff*, but introduction of split sites could affect *koff* of sFRB pairs. To test for irreversibility, cells expressing PM-localized Lyn-YFP-sFRBχ_N_, cytosolic CFP-sFRBχ_C_ (χ = 1, 2, 3) and cytosolic mCh-FKBP were incubated with rapamycin for 30 mins. Samples were then washed ten times, and cells showing trimerization were imaged over 30 mins (**Supplementary Fig. 5a**). No appreciable decrease in PM/cytosol ratio was detected for either CFP-sFRBχ or mCh-FKBP (**Supplementary Fig. 5b**), indicating that sFRB CIT systems are practically irreversible. To investigate whether the three sFRB pairs work in an orthogonal manner, we alternated sFRB_N_ with sFRB_C_ halves split at different sites. While we determined that sFRB pairs are non-orthogonal, we discovered three additional sFRB pairs (sFRB_1N_/sFRB_3C_, sFRB_2N_/sFRB_1C_, sFRB_2N_/sFRB_3C_) that trimerize with FKBP (**Supplementary Fig. 6**).

### Structural validation of split FKBP

First, we solved the crystal structure of a complex of unsplit FKBP12 and FRB (T2098L) proteins with rapamycin to examine whether the point mutation of FRB affects the overall conformation and/or the interaction with rapamycin. The crystal structure of FKBP12 · rapamycin · FRB (T2098L) determined at 2.20 Å resolution (**Fig. 3a**) shows only 0.61 Å of the average root-mean-square deviation (RMSD) for Cα atoms to the wild-type FRB-containing complex (PDB ID: 1FAP), indicating that the both structures are in almost perfect agreement. Also, the major interactions between FRB (T2098L), FKBP12 and rapamycin were the same as in the wild-type FRB · FKBP12 · rapamycin complex (**Fig. 3b**). Hence, we confirmed that the T2098L mutation of FRB does not affect the overall conformation and the interactions with rapamycin.

Next, we attempted to purify sFRBχ and sFKBPχ fragments and reconstitute their complexes to structurally validate our split strategy. We could purify sFKBP_1N_ as an MBP fusion protein by an *E. coli* expression system although we could not obtain the sFRBχ_N_ fragments as a soluble protein. Therefore, we aimed to structurally validate whether sFKBP_1N_ and sFKBP_1C_ form a functional sFKBP_1_ complex. We solved crystal structures of sFKBP_1_·rapamycin and sFKBP_1_·rapamycin·FRB (T2098L) at 2.92–3.11 Å resolution (**Fig. 3c,d**). In both structures, sFKBP_1N_ and sFKBP_1C_ formed the same secondary structure as FKBP12, showing only 0.38 and 0.42 Å of the average RMSD for Cα atoms to the corresponding region of the intact FKBP12 protein, respectively. The interactions between sFKBP_1_ and rapamycin are completely conserved in both structures. Finally, the overall structure of sFKBP_1_·rapamycin · FRB (T2098L) shows only 0.79 Å of the average RMSD to the intact FKBP12 · rapamycin · FRB (T2098L) complex, without disrupting hydrogen bonds and major hydrophobic interactions to FRB and rapamycin (**Fig. 3e**). These results indicate that, *in vitro*, the split FKBP fragments form a functional FKBP protein which is structurally indistinguishable to the intact FKBP protein.

### CIT at organelle membrane junction sites

Membrane contact sites (MCS), also called organelle-organelle junctions, are subcellular regions of close inter-organellar membrane apposition where unique signaling can occur, allowing organelles of specialized biochemical function to coordinate with each other while retaining their distinct identities^4,19,20^· MCS are widespread among almost all membrane-bound organelles^21,22^, and local MCS molecular tethers and actuators^23^ regulate processes ranging from calcium^24,25,26^ and lipid transport^27^ to organelle fission^28,29^. Functional studies for MCS are complex; most MCS regulate multiple functions, and local MCS proteins play compensatory roles, such that few MCS have been completely ablated by the knock-down or knock-out of a single gene^30^. These confounding conditions make rapidly inducible perturbation attractive for studying MCS, as supported by prior CID-based studies in the field^4, 19, 20, 31^. As proof-of-principle, we used CIT to target cytosolic protein to ER-mitochondria and ER-PM MCS, demonstrating its potential as a generalizable recruitment-based screen for MCS function (**Fig. 4a**).

**Figure 4:**
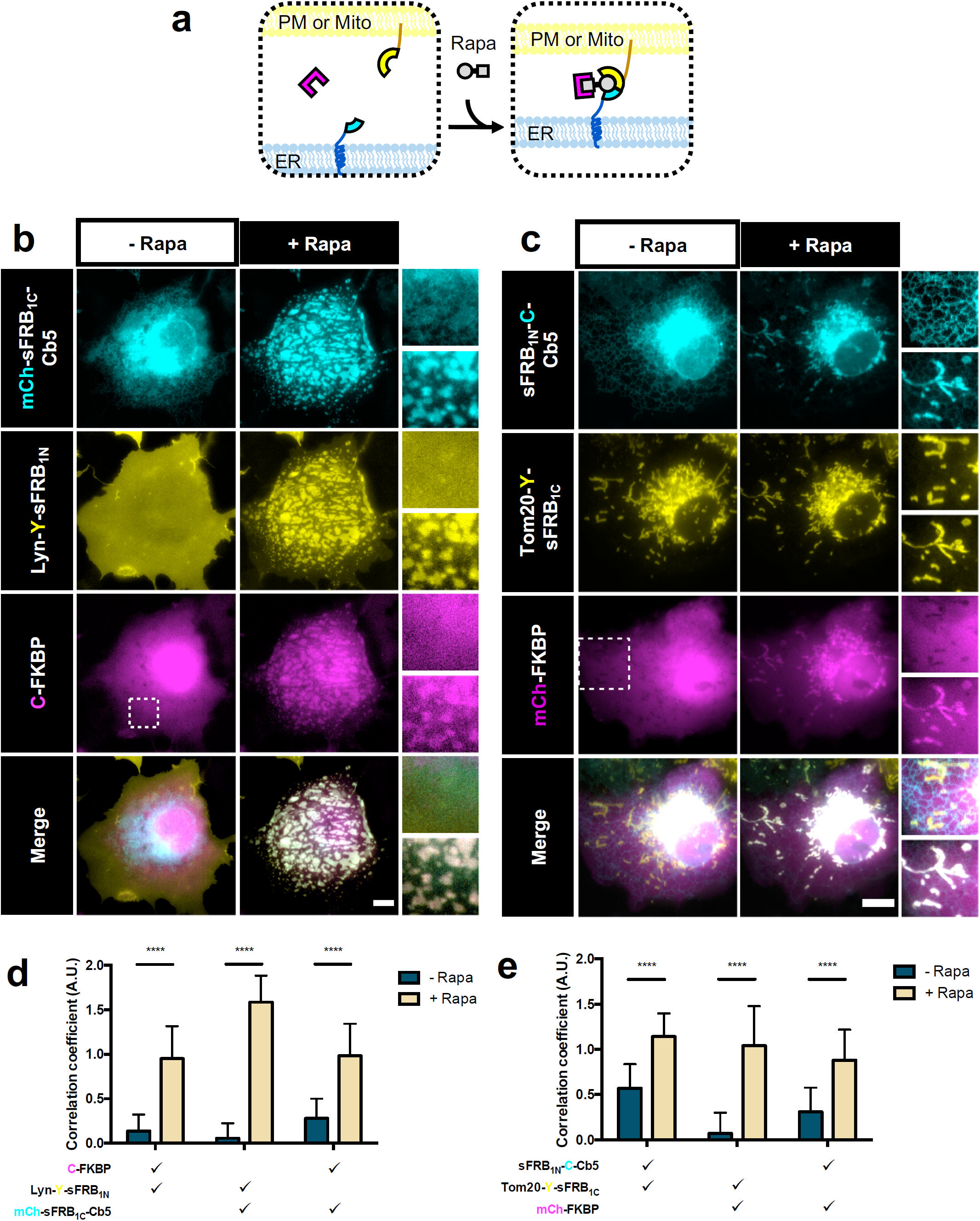
CIT-based recruitment of cytosolic protein to ER-PM and ER-mitochondria MCS. (a) Targeting cytosolic FKBP (magenta) to ER-PM or ER-mitochondria MCS. sFRB halves (cyan and yellow) targeted to two organelles recruit FKBP in a rapamycin-dependent manner. (b) Cos-7 cells expressing mCh-sFRB_1C_-Cb5, Lyn-Y-sFRB_1N_ and CF pre- and 12 min post-rapamycin for CF recruitment at ER-PM MCS. (c) Cos-7 cells expressing sFRB_1N_-C-Cb5, Tom20-Y-sFRB_1C_, and CF pre- and 12 min post-rapamycin for CF recruitment at ER-mitochondria MCS. (d, e) Quantifying trimerization between the 3 signals pre- and post-rapamycin. Check marks specify each combination of two wavelengths used in calculating pairwise Fisher’s transformation of Pearson’s correlation coefficients. Student’s t-test was used to compare correlations pre- and post-rapamycin (d, e: n = 38, 27; 3 experiments). Scale bars, 10 μm.

To recruit cytosolic protein to ER-PM junctions, we co-expressed PM-targeted sFRB_1N_ (Lyn-CFP-sFRB_1N_) and ER-targeted sFRB_1C_ (YFP-sFRB_1C_-Cb5) with cytosolic mCh-FKBP (**Fig. 4a**). Upon rapamycin addition, ER and PM tethering formed patches that colocalize with mCh-FKBP (**Fig. 4b** and **Supplementary Video 1**). Pairwise image correlation between all three image wavelengths all significantly increased after rapamycin (p <0.0001, **Fig. 4d**), suggesting that the CIT system can recruit cytosolic protein to the ER-PM MCS. When we removed sFRB_1N_ (using Lyn-CFP only) or sFRB_1C_ (using only YFP-Cb5) from the experiment, trimerization could not be observed (**Supplementary Fig S7a,b**, left and center) with no increase in image correlation coefficients between all three signals. However, when mCh alone was expressed instead of mCh-FKBP, ER-PM tethering still occurred, albeit with no mCh recruitment to the site, and the image correlation coefficient between the ER and PM signals significantly increased (**Supplementary Fig S7a,b**, right).

We also used sFKBP to recruit cytosolic FRB-mCh to ER-PM junctions. Cells showed undetectable or prominent FRB-mCh recruitment at ER-PM junctions pre- and 1 h post-rapamycin addition (**Supplementary Fig. 8a,b**, respectively). Imaging showed much slower cytosolic protein recruitment compared to sFRB trimerization (12 mins vs. 1 h, **Supplementary Video 1** vs. **Supplementary Video 2**). Pairwise image correlation of the three wavelengths showed significantly increased correlation coefficient (p<0.0001) between the ER and PM channels, increased correlation between cytosolic FRB-mCh and ER (p=0.030), and increased but not significant cytosolic FRB-mCh and PM correlation (p=0.16), reflecting less robust FRB-mCh recruitment (**Supplementary Fig. 8c**). Because sFKBP was less efficient compared to sFRB in targeting cytosolic proteins to ER-PM junctions, we decided to continue with only sFRB for MCS applications for the rest of the study.

We applied sFRB CIT further to recruit cytosolic proteins to ER-mitochondria junctions. Complimentary sFRB pairs were expressed on the ER (sFRB_1N_-CFP-Cb5) and mitochondria (Tom20-YFP-sFRB_1C_), and post-rapamycin addition translocated mCh-FKBP to ER-mitochondria MCS (**Fig. 4c** and **Supplementary Video 3**). Pairwise image correlation coefficients increased for all three wavelengths, indicating successful trimerization (p <0.0001 all cases; **Fig. 4e**). Likewise, removal of sFRB_1N_ (Lyn-YFP) and sFRB_1C_ (mCh-Cb5) yielded no translocation of mCh-FKBP to ER-mitochondria MCS after rapamycin addition (**Supplementary Fig. 9a,b**, left and center), whereas removal of FKBP resulted in increased co-localization between the ER and mitochondria components, without any co-localization of mCh (**Supplementary Fig. 9a,b**, right).

### CIT induces tri-organellar membrane contact sites

Multi-spectral live-cell imaging of organelles has revealed that ER-mitochondria MCS make frequent contact with the Golgi, peroxisomes, and lipid droplets, hinting at tri-organellar MCS^22^. ER-mitochondria MCS also mark regions of autophagosome formation^32^. We therefore speculated that a tri-organelle ER-PM-mitochondria junction site could exist due to the existence of ER-PM, ER-mitochondria, and PM-mitochondria MCS, and their shared function in Ca^2+^ signaling. Using CIT to determine if a tri-organellar junction could be synthetically formed between the PM, ER and mitochondria, we co-expressed sFRB_1C_ localized to the PM (Lyn-CFP-sFRB_1C_), sFRB_1N_ to the ER (YFP-sFRB_1N_-Cb5) and FKBP to the mitochondria (Tom20-mCh-FKBP) in Cos-7 cells (**Fig. 5a-c**). Fifteen minutes post-rapamycin addition, PM, ER and mitochondrial signals colocalized (**Fig. 5b,c** and **Supplementary Video 4**); correlation coefficients increased between PM-ER, ER-mitochondria and PM-mitochondria signals (p < 0.0001 all cases; **Fig. 5d**). When sFRB_1C_, sFRB_1N_, or FKBP were each removed, trimerization no longer occurred post-rapamycin (**Supplementary Fig. 10a,b**, left and right), but removal of FKBP still resulted in colocalization of ER-PM signals (**Supplementary Fig. 10a,b**, center). These results show that CIT can induce PM-ER-mitochondria tethering, and serves as a detection method for potential tri-organellar MCS formation between these three organelles.

**Figure 5:**
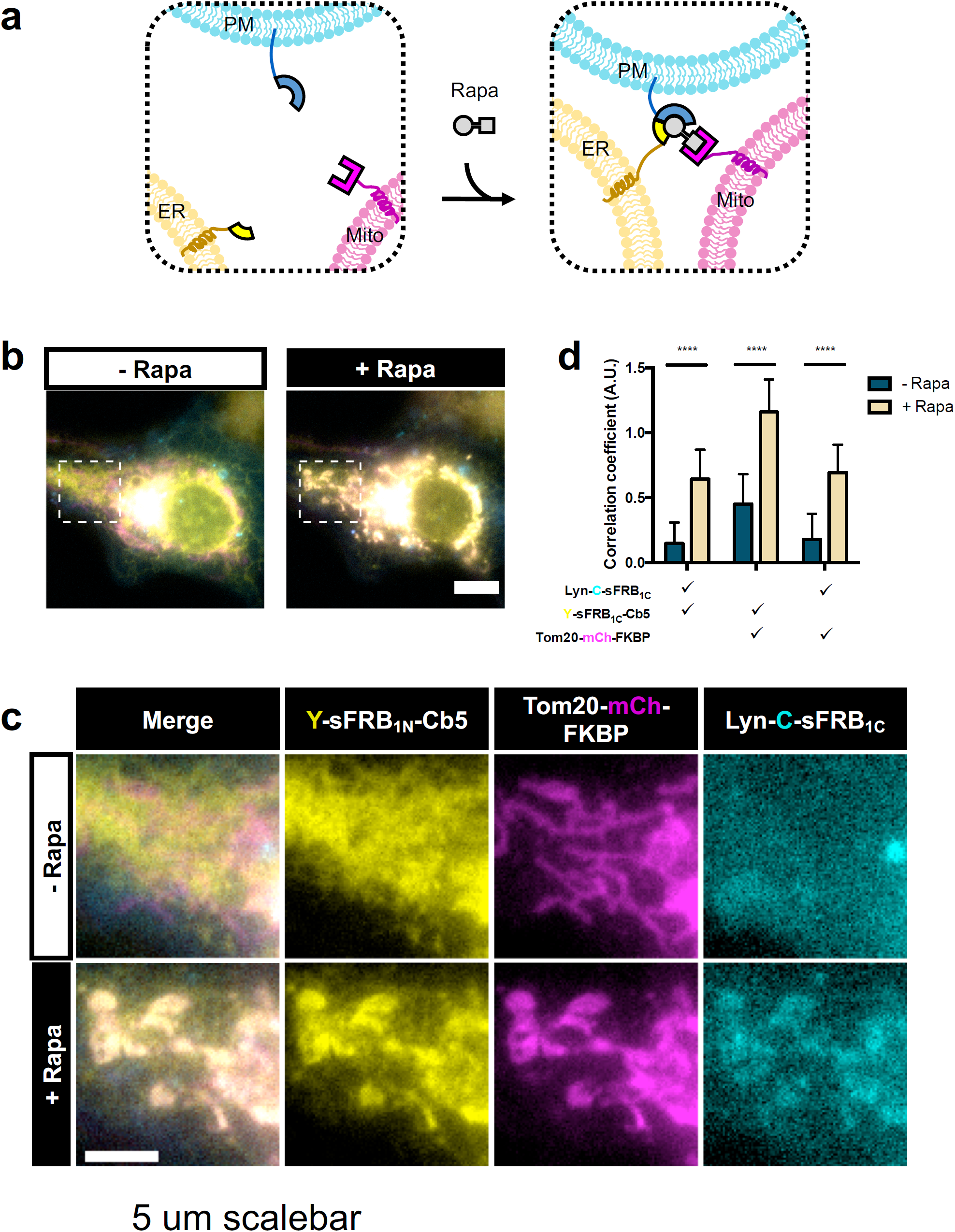
Tri-organellar junction formation by CIT. (a) Trimerization of CIT components targeted to ER, mitochondria and PM by rapamycin. (b, c) Cos-7 cell expressing Y-sFRB_1N_-Cb5, Tom20-mCh-FKBP and Lyn-C-sFRB_1C_ pre- and 15 mins post-rapamycin. (d) Quantification of trimerization extent through pairwise correlation of Fisher’s transformation of Pearson’s correlation coefficient between three wavelengths (*n* = 23 cells; 4 experiments). Scale bars, 5 μm.

### Localized PI(4,5)P_2_ depletion at ER-PM junctions

The major functions of ER-PM MCS are Ca^2+^ re-entry and lipid homeostasis. As Ca^2+^ re-entry is more established, the focus in the ER-PM field has shifted in recent years towards lipid transfer and homeostasis^33^. One major topic in the field is the regulation of secondary lipid messenger PI(4,5)P_2_ (PIP_2_) and its precursor PI(4)P. Because the PM is a large signaling hub of the cell converting messages from extracellular milieu to intracellular signals, PIP_2_ and its precursor PI(4)P levels are often in flux. ER-PM junctions serve as a bridge connecting the PM and ER, where local proteins regulate PM lipid homeostasis by using the ER as a reservoir or a sink, depending on the needs of the PM^20, 31, 34^. While methods exist to perturb PM lipid composition or locally deplete PIP_2_ from the PM^35^, the means of perturbing lipids only at ER-PM junctions have yet to be established.

To achieve local PIP_2_ depletion, we targeted cytosolic PIP_2_ phosphatase, INP54P, to ER-PM junctions. PM-localized sFRB_1N_ (Lyn-Clover-FRB_1N_), ER-localized sFRB_1C_ (iRFP-FRB_1C_-Cb5), cytosolic CFP-FKBP fused to 5-phosphatase INP54P (331) (CF-INP54P), and mRuby-PH-PLCδ, a biosensor for PIP_2_ were co-expressed in Cos-7 cells (**Fig. 6a**). After rapamycin addition, CF-INP54P was recruited to ER-PM junctions, and a marked pattern of reduction in mRuby-PH-PLCδ intensity was observed (**Fig.6b**, left, and **Supplementary Video 5**). In contrast, recruitment of the INP54P phosphatase-dead mutant CF-INP54P D281A, or CF alone did not reduce mRuby-PH-PLCδ intensity (**Fig.6b**, center and left, and **Supplementary Videos 6 and 7**, respectively). Localized PIP_2_ reduction occurred in areas of high (arrow) compared to low ER-PM junction formation (chevron), reflected in percent change in mRuby-PH-PLCδ intensity in the respective regions (p < 0.0001, **Fig. 6c**). CF-INP54P D281A recruitment inside ER-PM rich regions was slightly decreased compared to outside (p = 0.0358, **Fig. 6c**). CF recruitment inside ER-PM rich regions was not significantly different compared to outside (p = 0.869, **Fig. 6c**). These results show that CIT-based recruitment of INP54P can locally deplete PIP_2_ at ER-PM junctions.

**Figure 6:**
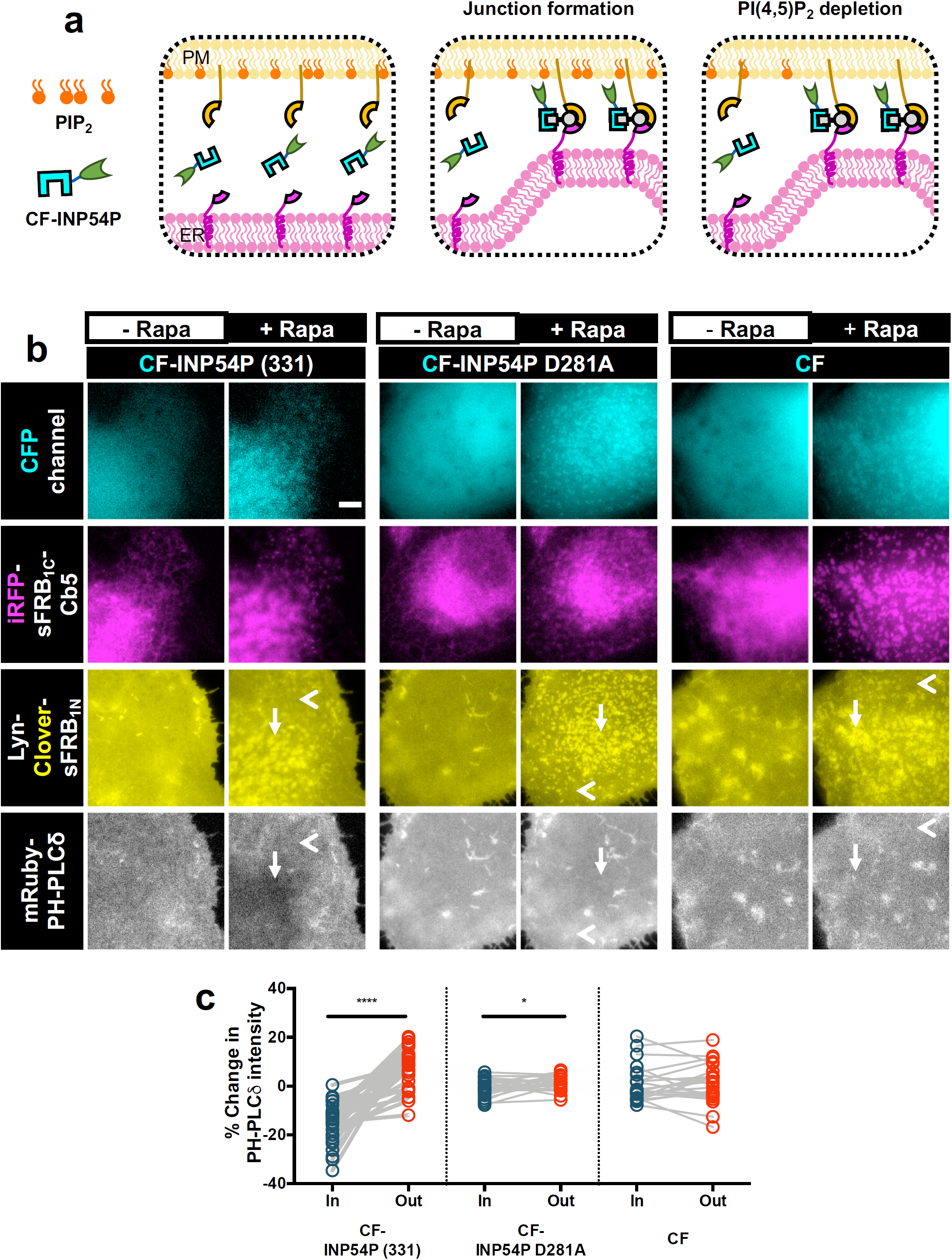
CIT induces local PI(4,5)P2 depletion at ER-PM MCS. (a) Schematic of CIT-induced PI(4,5)_2_ depletion. CF-INP54P is recruited to ER-PM MCS by ER and PM localized sFRB halves. Upon junction formation (middle panel), INP54P accesses PM PI(4,5)P_2_ leading to local PI(4,5)P_2_ depletion (right panel). (b) Comparing effects of ER-PM MCS recruitment of functional CF-INP54P (331) against phosphatase-dead mutant CF-INP54P (D281A) and CF alone. Cos-7 cells also express Lyn-Clover-sFRB_1N_, iRFP-sFRB_1C_-Cb5 and mRuby PH-PLCδ, a PI(4,5)P_2_ biosensor. Time-lapse epifluorescence microscopy taken pre- and 5 mins post rapamycin addition. (c) Intensity of mRuby PH-PLCδ PI(4,5)P_2_ biosensor taken from regions of inside (arrow) and outside (chevron) ER-PM MCS, significance analyzed with paired Student’s t-test (from left to right: *n* = 28, 25 and 23 cells; each 4 experiments). Scale bar, 10 μm.

## Discussion

In this study we designed, developed and implemented the first known CIT system by rationally splitting proteins of the FKBP/FRB CID system. The CIT tool is small, robust, and trimerizes on a timescale of seconds-to-minutes. We used CIT to rapidly target cytosolic protein exclusively to ER-PM and ER-mitochondria MCSs, induce tri-organellar ER-mitochondria-PM junctions and locally deplete PIP_2_ at ER-PM MCS. We believe that our ability to achieve CIT is attributed to the unique cooperativity of the FKBP/FRB CID. This efficient CIT system would be otherwise highly difficult to achieve *de novo*, as engineering sequential cooperativity at protein interfaces would be necessary.

The sFKBP CIT system, being slower than the sFRB CIT system, provides evidence for the mechanism of trimerization. As FKBP binding affinity for rapamycin is over 10,000-fold higher than FRB, FKBP-rapamycin complexes form much faster. We believe that in sFRB systems, FKBP-rapamycin complexes are first quickly formed, then bind and stabilize sFRB halves, resulting in trimerization. In contrast, in sFKBP systems, FRB-rapamycin complexes take longer to form, causing slow stabilization of the trimer. It is also possible for split proteins to bind free rapamycin first, and then recruit its CID partner; indeed, sFKBP can dimerize *in vitro* in the absence of FRB, as evidenced by crystal structure analysis (**Fig. 3**). However, it is relatively unlikely to expect split proteins to bind rapamycin with higher affinity than its full length counterpart. Therefore our results suggest that the predominant mechanism of trimerization is split protein recombination mediated by the rapamycin-CID partner complex.

Components of sFRB and sFKBP CIT systems are small; sFRB_1N_ and sFRB_1C_ are 5 kDa and 6.3 kDa respectively, thereby increasing functionality without adding bulk. While sFRB_χN_ and sFRB_χC_ halves do not function orthogonally, we discovered three additional sFRB pairs (sFRB_1N_/sFRB_3C_, sFRB_2N_/sFRB_1C_, sFRB_2N_/sFRB_3C_) that trimerize with FKBP (**Supplementary Fig. 6d**). Thus in total, we have developed 6 sFRB pairs capable of trimerization, increasing the likelihood of successful CIT implementation across biological systems. Having 6 CIT systems introduces variety for future mutagenesis studies to achieve orthogonality, irreversibility and improved expression.

Depletion of PIP_2_ at ER-PM junctions (**Fig. 6**) serves as a springboard for further experiments investigating differences in local PIP_2_ depletion at the ER-PM junction compared to global PIP_2_ depletion at the PM; it may provide insight into the relationship between lipid transfer and Ca^2+^ entry processes at the ER-PM junction, as they share common players such as PIP_2_ and PI(4)P. In a broader context, CIT-based targeting of panels of molecular actuators or proteins of interest to junctions can serve as a top-down screening method to investigate local signaling. Additionally, the tri-organellar PM-ER-mitochondria junction formed through CIT indirectly suggests that these three organelles have the potential of interacting. In the future, CIT can be a powerful tool to validate and perturb other tri-organellar MCS, such as the ER-mitochondria-Golgi junction suggested by a recent study^22^, which is feasible to achieve due to an inherently modular nature of CIT molecular design.

In all, we developed a trimerization tool capable of cell signaling manipulation in new ways. MCS targeting in this study demonstrated the enhanced spatiotemporal specificity CIT has over CID. We see potential for the tool beyond its role in future screens for studying MCS. Trimerization is a common mechanism in nature, and CIT-based mimicry may deduce the significance of trimerization in these systems. CIT may also produce increasingly sophisticated, tunable switches and logic gates. By introducing a third component to be recruited to two others with one chemical stimulus, CIT introduces a new tool to cell, chemical and synthetic biology toolkits.

## Online Methods

### Cell culture and transfection

HeLa and Cos-7 cells were maintained at 37°C, 5% CO2 and 95% humidity in Dulbecco’s modified Eagle’s medium (DMEM) supplemented with 10% FBS and 1% Pen/strep (Gibco). The day prior to imaging, cells were detached with 5% Trypsin-EDTA (Gibco) and harvested for reverse transfection with FuGENE HD (Promega). As a general rule, co-transfection of split and unsplit proteins used a split protein to unsplit protein plasmid ratio of 20:1 for sFRB CIT experiments and a ratio of 10:1 for sFKBP CIT. Prior to imaging, cell medium in samples was replaced with DPBS containing Ca^2+^ and Mg^2+^ for imaging < 30 mins or fresh DMEM growth media for imaging > 30 mins. Cells were transfected with constructs at adjusted relative plasmid ratios and incubated for 16-24h prior to imaging.

### DNA plasmid construction

Constructs used to determine working split proteins were based off Clontech C1 and N1 backbones with various fluorophores. A 5xSAGG flexible linker oligo was annealed was inserted between KpnI and SalI restriction sites of both C1 and N1 backbones. Lyn or CAAX oligo DNA fragments were subsequently annealed and ligated into C1 or N1 backbones containing 5xSAGG linkers. Then, FRB or FKBP fragments were cloned and inserted between KpnI and BamHI or HindIII and SalI for C1 and N1 vectors respectively. To obtain non-Lyn or CAAX versions of the proteins, the portion containing fluorescent protein with split FRB or FKBP fragments were cut and pasted into C1 or N1 backbone vectors lacking Lyn or CAAX. YFP-sFRB_1N_-Cb5 was generated by deletion mutagenesis of previously reported YFP-FRB-Cb5^4^. ER-targeting sequence Cb5 was amplified by PCR and subcloned into a sFRB_1N_-C backbone from sFRB_1N_-C-Cb5 to generate sFRB_1N_-C-Cb5. CF-INP54P (331) and CF-INP54P (D281A) were from a previous report 36. All plasmids were verified by Sanger sequencing.

### Live-cell imaging

Most imaging was conducted on a IX71 microscope (Olympus) with a 63x objective (and 1.6x zoom) and an ORCA-Flash4.0 LT Digital CMOS camera (Hamamatsu). Rate constant of FKBP recruitment by sFRB_1_ was imaged at 10 s intervals with a spinning disc confocal, inverted Axiovert 200 (Zeiss) with a 40x objective and an Orca ER CCD camera (Hamamatsu). Both microscopes were driven by Metamorph 7.5 imaging software (Molecular Devices). Four-color imaging in Fig. 4 was conducted with an Eclipse Ti microscope (Nikon) with a 60x objective (and 1.5x zoom) and a Zyla 4.2 sCMOS camera (Andor), driven by NIS Elements software (Nikon). Unless otherwise indicated imaging was done at 1-3 min intervals for 12-30 mins, at times with between 3-5 0.5 um-spaced z positions. Images analyzed and shown are from a single plane, and not maximum intensity projections. Microscopy experiments applying CIT to membrane contact sites were conducted at 37°C, 5% CO_2_ and humidity with a stage top incubation system (Tokai Hit).

### Analysis of PM-to-cytosol ratio

PM-to-cytosol ratio was determined through linescan analysis. Lines through cells were drawn to 1) avoid the nucleus and 2) maximize the two membrane signals on both sides of the cell. Two PM (p1, p2) locations based on peak intensities (I) along the linescan were determined visually. The cytosol was defined as midpoint location (m) between p_1_ and p_2_. A moving average of 3 was used to determine signal intensity at p_1_ and p_2_, and a moving average of 5 at the cytosol location. Then PM-to-cytosol ratio was determined for each PM location, and then averaged. Percent change was determined from PM to cytosol ratio values before and after rapamycin treatment.

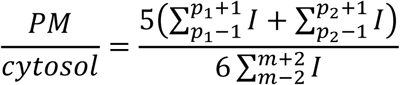

### Analysis of recruitment rate

Averaged intensities from 3 regions of interest (ROI’s) from the cytosol were used to generate exponential decay curves. Averaged intensity timepoints with exponential decay trends were selected for line fitting with an exponential regression to determine the exponential recruitment coefficient. R^2^ values were all > 0.88.

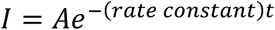

### Analysis of trimerization with Fisher’s transform of Pearson’s correlation coefficient

3 ROI’s about 50 x 50 pixels were manually positioned for each cell to determine Pearson’s correlation using FIJI. ROIs were chosen include enough background and organelle to mitigate false correlations. Due to organelle movement and deformation, different ROI’s were selected for the same cell before and after rapamycin addition. Pairwise Pearson’s correlation (between ROI1 and ROI2) was determined among three channels using the built-in MATLAB function corr2, and averaged between 3 ROI’s. Fisher’s transform was performed to normalize Pearson’s correlation coefficient for later statistical analysis. The result is altogether referred to as correlation coefficient.

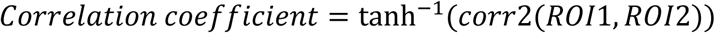

### Statistical analysis and data visualization

Unless otherwise stated, most statistical analysis was done with unpaired, two-tailed Student’s T-test assuming equal distributions. A paired, two-tailed Student’s T-test was used in Fig. 6c. PRISM 6 software was used for statistical analysis and data visualization.

### Split site prediction

Split sites were determined by selecting non-evolutionarily conserved solvent accessible loops. The Stride program ^37^ was used to calculate the solvent accessible areas (SAA). Loops exceeding 30 Å^2^ were selected as potential split sites. Sequence conservation as indicated by Kullback-Liebler conservation scores was determined with MISTIC ^38^ and FKBP domain and FRB domain sequences used were obtained from PFAM server ^39^ (Finn 2014). Loops with Kullback-Leibler conservation scores less than 2 were selected as potential split sites. The difference between the native energy of the intact protein and the sum of the energies of the split proteins at a given split site - the split energy - was calculated using the MEDUSA scoring function ^40^ (Yin 2007). Energy wells were indicative of split sites in this study, which differs from the previous implementation of SPELL, which targeted regions outside energy wells ^41^. The original SPELL study looked for split proteins sites which minimized the destabilization as these proteins were then fused to a known destabilizer domain (iFKBP) ^41^. This approach ensures the prevention of the split part association in the absence of rapamycin.

### Crystallography

#### Protein expression and purification

For structural analysis, cDNA encoding the sFKBP_1N_ (residues 1–32 of human FKBP12) was amplified by PCR and subcloned into a pET28a(+) derivative vector, which has a maltose binding protein (MBP)-encoding sequence inserted after an N-terminal polyhistidine tag. The construct contained a tobacco etch virus (TEV) protease recognition site at the junction between polyhistidine tag and the MBP sequences. Recombinant sFKBP_1N_ protein was expressed in *E. coli* BL21 (DE3) cells in LB broth at 37 °C until the OD_600_ reached 0.8, at which point the temperature was shifted to 18 °C. Protein expression was induced by addition of isopropyl-β-D-thiogalactopyranoside to a final concentration of 250 µM. The cultures were grown for an additional 20 hrs and harvested by centrifugation. Cell pellets were resuspended in 50 mM Tris-HCl buffer (pH 7.0) containing 500 mM NaCl, 20 mM imidazole and 10% glycerol. Cells were lysed by sonication and clarified by centrifugation. The cell lysate was loaded onto a HisTrap HP column (GE Healthcare), and eluted with 50 mM Tris-HCl buffer (pH 7.0) containing 500 mM NaCl, 500 mM imidazole and 10% glycerol. The N-terminal histidine tag was cleaved by incubating with TEV protease overnight at 4 °C. The protein was further purified using a HiLoad Superdex 200 16/60 size-exclusion column (GE Healthcare) equilibrated with 20 mM Tris-HCl buffer (pH 8.0) containing 150 mM NaCl and 5 mM DTT. The sFKBP_1C_ (residues 33–108 of human FKBP12) was expressed and purified essentially as the same as sFKBP_1N_, except an N-terminal His-SUMO tag was fused instead of the polyhistidine and MBP. The His-SUMO tag was cleaved by SUMO protease overnight at 4 °C. The purified sFKBP_1N_ and sFKBP_1C_ proteins were mixed in a 1:2 molar ratio, and then the mixture was added to a 1 µM rapamycin solution in 20 mM Tris-HCl buffer (pH 8.0) containing 150 mM NaCl, 5 mM DTT and 10% DMSO. The sFKBP_1N_, sFKBP_1c_ and rapamycin mixture was filtered and loaded onto a Superdex 200 Increase 10/300 column (GE Healthcare) equilibrated with 20 mM Tris-HCl buffer (pH 8.0) containing 150 mM NaCl and 5 mM DTT. The eluted peak fractions of sFKBP_1_·rapamycin complex were pooled and concentrated to 40 mg/ml. The cDNAs encoding human FKBP12 (residues 1–108) and FKBP12·rapamycin binding (FRB) domain encompassing residues 2021–2113 of human mTOR (T2098L) were subcloned into pET28a(+) containing an N-terminal polyhistidine-GST tag. The FKBP12 and FRB mutant proteins were expressed and purified essentially as the same as sFKBP_1N_. The His-GST tag was cleaved by TEV protease overnight at 4 °C.

#### Crystallization, data collection and structure determination

Crystallization was performed by the sitting-drop vapor-diffusion method at 20 °C. Crystals of sFKBP_1_ · rapamycin appeared by mixing equal volumes of a 40 mg/ml sFKBP_1_ · rapamycin complex solution and a reservoir solution of 100 mM Na-HEPES buffer (pH 7.5) containing 20% (w/v) PEG 8000. FKBP12 · rapamycin · FRB (T2098L) complex was prepared by mixing them at an equimolar ratio and incubating for 60 min on ice. Crystals of FKBP12 · rapamycin · FRB (T2098L) appeared at 100 mM cacodylic acid buffer (pH 6.5) containing 350 mM zinc acetate and 8% (w/v) isopropanol. sFKBP_1_ · rapamycin·FRB (T2098L) complex was prepared in the same manner as FKBP12 · rapamycin·FRB (T2098L). Crystals of sFKBP_1_ · rapamycin · FRB (T2098L) appeared at 100 mM Tris-HCl buffer (pH 7.0) containing 200 mM calcium acetate and 20% (w/v) PEG 3000. All crystals were briefly soaked in a cryoprotectant drop composed of the reservoir solution supplemented with 20% glycerol and then flash-cooled in liquid nitrogen for X-ray diffraction data collection. The datasets were collected at 1.000 Å, 100 K on the beamline BL26B2 at SPring-8 (Harima, Japan) or the beamline X06DA at the Swiss Light Source (Villigen, Switzerland). They were processed using the *XDS* ^42^ and scaled using *AIMLESS* supported by other programs of the *CCP4* suite ^43^. Crystal structures were determined by molecular replacement using *MOLREP* ^44^ with the FKBP12·rapamycin·FRB structure (PDB ID: 1FAP) and the MBP structure (PDB ID: 1ANF) as the search models. Model building was accomplished with *Coot* ^45^, and structural refinement was performed with *REFMAC* ^46^ and *PHENIX* ^47^. The Ramachandran statistics are as follows: 96.0% favored, 4.0% allowed for sFKBP_1_ · rapamycin; 97.2% favored, 2.8% allowed for FKBP12·rapamycin · FRB (T2098L); and 96.8% favored, 3.2% allowed for sFKBP_1_·rapamycin · FRB (T2098L). The data collection and refinement statistics are summarized in **Supplementary Table 1**. The structural models in the figures were depicted using *PyMOL* version 1.8 software (Schrödinger, LLC). Two-dimensional interaction plots were carried out with LIGPLOT ^48^.

## Acknowledgements

We thank Luca Bertozzi and Sarah Thompson for help with plasmid generation. We thank Robert DeRose, Xinyi Y. Zhou, and Yuta Nihongaki for proofreading the manuscript. We thank Hideaki Niwa (RIKEN), Naoki Sakai (RIKEN), and the staff at the BL26B2 beamline (Proposal No. 20190047) of SPring-8 (Harima, Japan) and the X06DA beamline (Proposal No. 20171001) of the Swiss Light Source, Paul Scherrer Institut (Villigen, Switzerland) for their help in X-ray diffraction data collection. We acknowledge support from the National Institutes for Health (5R01GM123130 to T.I., and 5R01GM123247 and 1R35 GM134864 to N.V.D.), the Passan Foundation to N.V.D., the DoD DARPA (HR0011-16-C-0139 to T.I.), and the PRESTO program of the Japan Science and Technology Agency to T.U. (No. JPMJPR12A3) and T.I (No. JPMJPR12A5), and a Grant-in-Aid for Scientific Research (B) to T.U. (No. 16H05089) from the Japan Society for the Promotion of Science.

## Author contributions

H.D.W. and H.N. conceived the study with input from T.I. H.D.W. carried out cell experiments and conducted image analysis with help from A.K.A. M.K. purified and crystalized split proteins, and determined protein structure by X-ray crystallography. O.D. conducted rational split site analysis. T.I., H.N., T.U., and N.V.D. supervised the project. H.D.W. wrote the manuscript in consultation with T.I. and with input from M.K., T.U., O.D., and N.V.D.

