## Supplementary Information for "Rational design and implementation of a chemically inducible hetero-trimerization system"

**Contents:**

1. Legends for supplementary figures 1-10
2. Supplementary figures 1-10
3. Supplementary Table 1
4. Captions for supplementary movies 1-7

**Supplementary Figure 1:** Possible binding configurations of CID and CIT. (a) A small molecule dimerizer binds both proteins (yellow and magenta) in one of four configurations (red dotted box). (b) A small trimerizer binds all three proteins (yellow, magenta, and blue) in one of eight configurations (red dotted box).

**Supplementary Figure 2: FRB split sites and cytosolic sFRB expression check.** (a) Sequence alignment of FRB used in this study with mTOR FRB regions across species. Aligned though EMBL-EBI ClustalW and visualized with ESPript 3.0. Double arrowheads indicate split sites. (b) Visualization of sFRB N- and C-terminal halves (blue and red) across 3 split sites (PDB 1AUE). (c) Expression check of fluorophore-fused sFRB.

**Supplementary Figure 3: sFRB dimerization in the absence of FKBP overexpression.** (a) Epifluorescence snapshots of HeLa cells overexpressing N-terminal sFRB on the PM and its corresponding C-terminal sFRB in the cytosol, but not cytosolic FKBP, pre and 12 mins post 100 nM rapamycin addition. All three split sFRB pairs are tested. (b) PM/cytosolic ratios of C-terminal sFRB signal in cells pre- and 12 min post- rapamycin addition were determined through user-defined linescan analysis (from left to right: *n* = 31, 26, and 28; 3 experiments). Scale bar, 10 µm.

**Supplementary Figure 4: FKBP split site generation and trimerization characterization.** (a) Sequence alignment of FKBP12 used in this study with FKBP12 across species. Aligned though EMBL-EBI ClustalW and visualized with ESPript 3.0. Double arrowheads indicate split sites. (b) Translocation of mCh-FRB from cytosol to PM in HeLa cells co-expressing PM-targeted $sFKBP_1$ – $sFKBP_4$ pairs and full length FKBP. Pre-rapamycin; top 3 rows. 9 mins post-rapamycin; bottom row. Linescans show corresponding normalized intensity of FRB-mCh signal. Scale bar, 5 µm.

**Supplementary Figure 5: Effects of rapamycin washout on sFRB-FKBP trimer stability in cells.** (a) HeLa cells overexpressing PM localized Lyn-Y-$sFRB_{\chi N}$, cytosolic C-$sFRB_{\chi C}$, and cytolic mCh-FKBP, $\chi$ being split sites 1, 2, and 3, treated with 100 nM rapamycin for 30 mins and washed 10x prior to image capture at time 0 mins. Cells were imaged over 30 mins. PM/cytosolic ratios of mCh-FKBP (b) C-$sFRB_{\chi C}$ in cells pre- and 12 min post- rapamycin addition were determined through user-defined linescan analysis (for both graphs from left to right: *n* = 21, 13, and 29; 3 experiments). Scale bar, 5 µm.

**Supplementary Figure 6: Assessing cross-reactivity of sFRB pairs.** (a) 30 mins post 100 nM rapamycin treatment, Cos-7 cells expressing Tom20-mCh-FKBP at mitochondria and co-expressing all combinations of CFP-$sFRB_{\chi N}$ and YFP- $sFRB_{\chi C}$, where $\chi$ are different. (b,c) For each combination of sFRB, quantifying trimerization by calculating pairwise Fisher's transformation of Pearson's correlation coefficients between YFP-$sFRB_{\chi C}$ and Tom20-mCh-FKBP (b) or between CFP-$sFRB_{\chi N}$ and Tom20-mCh-FKBP (c) (for both graphs from left to right: *n* = 26, 30, 23, 25, 23, and 27 cells; 3 experiments). (d) Summary of sFRB pair functionality when split sites are varied. Scale bar, 5 µm.

**Supplementary Figure 7: Negative controls for cytosolic FKBP recruitment to ER-PM MCS.** (a) Assessing contributions of each CIT component in CFP-FKBP recruitment to ER-PM MCS; left, middle and center panels correspond to constructs lacking ER, PM, and cytosolic CIT components. (b) Quantifying trimerization between the 3 signals pre- and post-rapamycin. Check marks specify each combination of two wavelengths used in calculating pairwise Fisher's transformation of Pearson's correlation coefficients. Student's t-test was used for all correlations pre- and 12 mins post- 100 nM rapamycin (from left to right: *n* = 38, 27, and 32 cells; 3 experiments). Scale bar, 10 µm.

**Supplementary Figure 8: sFKBP-based CIT recruitment of cytosolic FRB to ER-PM MCS.** (a, b) Recruitment of mCh-FRB to ER-PM junctions by ER and PM targeted $sFKBP1_N$ and $sFKBP_{1C}$ pre- and 1 h post-rapamycin addition. FRB-mCh recruitment can be undetectable (a) or prominent (b). (c) Quantifying trimerization between the 3 signals pre- and post-rapamycin. Check marks specify each combination of two wavelengths used in calculating pairwise Fisher's transformation of Pearson's correlation coefficients. Student's t-test was used for all correlations pre- and post- 100 nM rapamycin (n = 24 cells; 3 experiments). Scale bar, 10 µm.

**Supplementary Figure 9: Negative controls for cytosolic FKBP recruitment to ER-mitochondria MCS.** (a) Assessing contributions of each CIT component in CFP-FKBP recruitment to ER-mitochondria MCS; left, middle and center panels correspond to constructs lacking ER, mitochondria, and cytosolic CIT components. (b) Quantifying trimerization between the 3 signals pre- and post-rapamycin. Check marks specify each combination of two wavelengths used in calculating pairwise Fisher's transformation of Pearson's correlation coefficients. Student's t-test was used for all correlations pre- and 12 mins post- 100 nM rapamycin (from left to right: *n* = 32, 26, and 24 cells; 3 experiments). Scale bar, 10 µm.

**Supplementary Figure 10: Negative controls for CIT-induced ER-mitochondria-PM tri-organellar membrane contact sites (MCS).** (a) Assessing contributions of each CIT component in tri-organellar MCS formation; left, middle and center panels correspond to constructs lacking ER, mitochondria, and PM CIT components. (b) Quantifying trimerization between the 3 signals pre- and post-rapamycin. Check marks specify each combination of two wavelengths used in calculating pairwise Fisher's transformation of Pearson's correlation coefficients. Student's t-test was used for all correlations pre- and 15 mins post- 100 nM rapamycin (from left to right: *n* = 24, 32 and 28 cells; 3 experiments). Scale bar, 10 µm.

Supplementary Figure 1

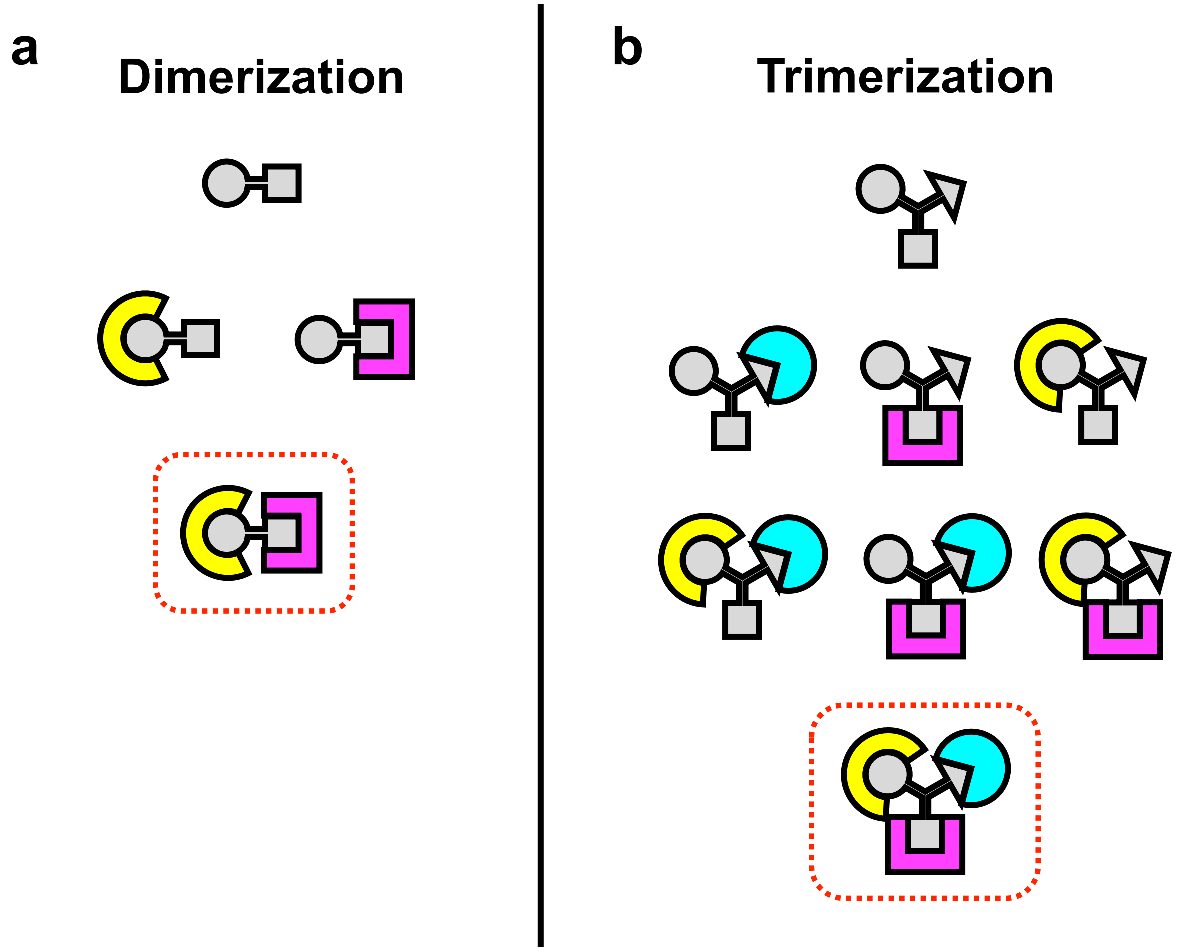

### Supplementary Figure 2

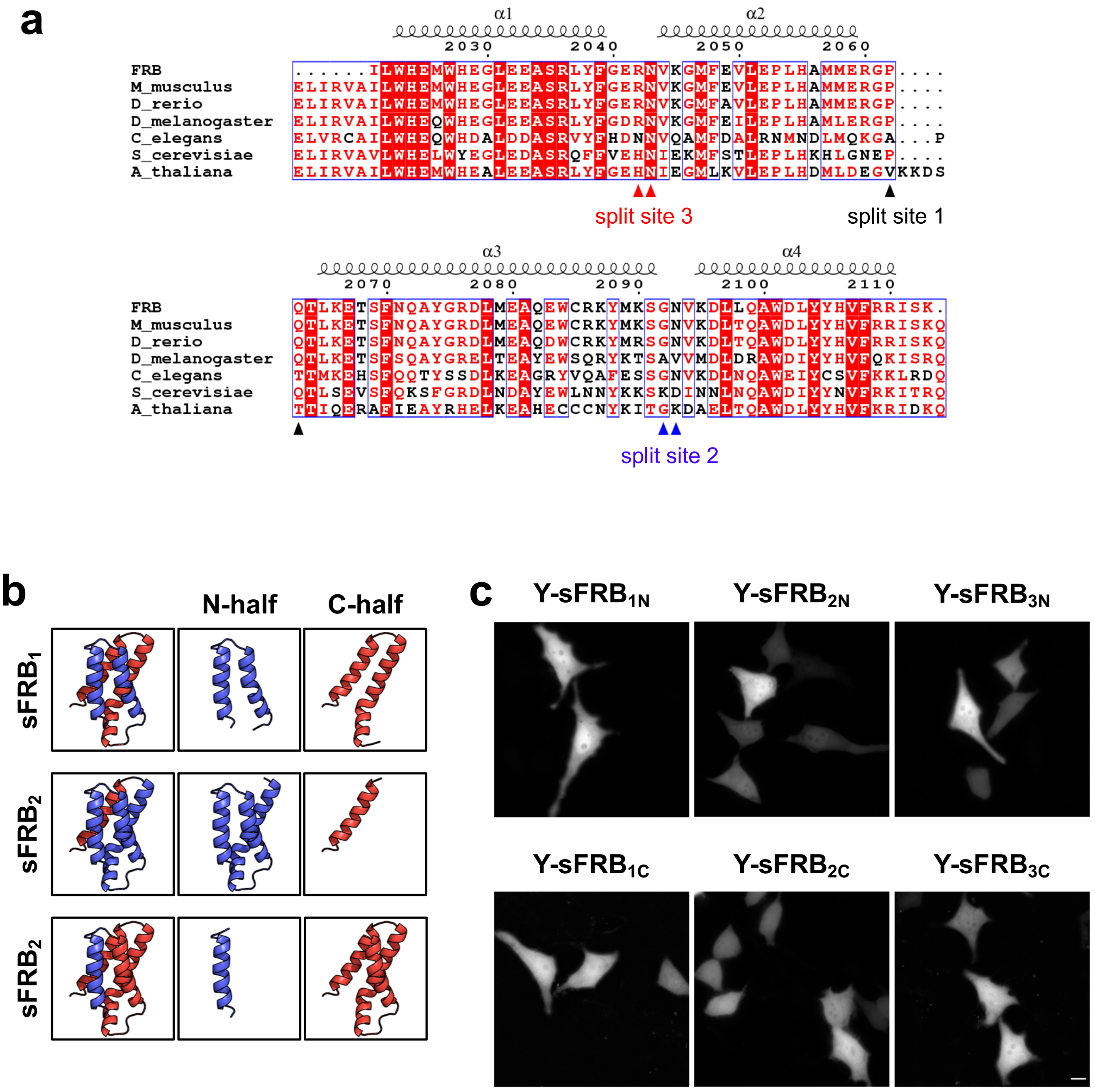

Supplementary Figure 3

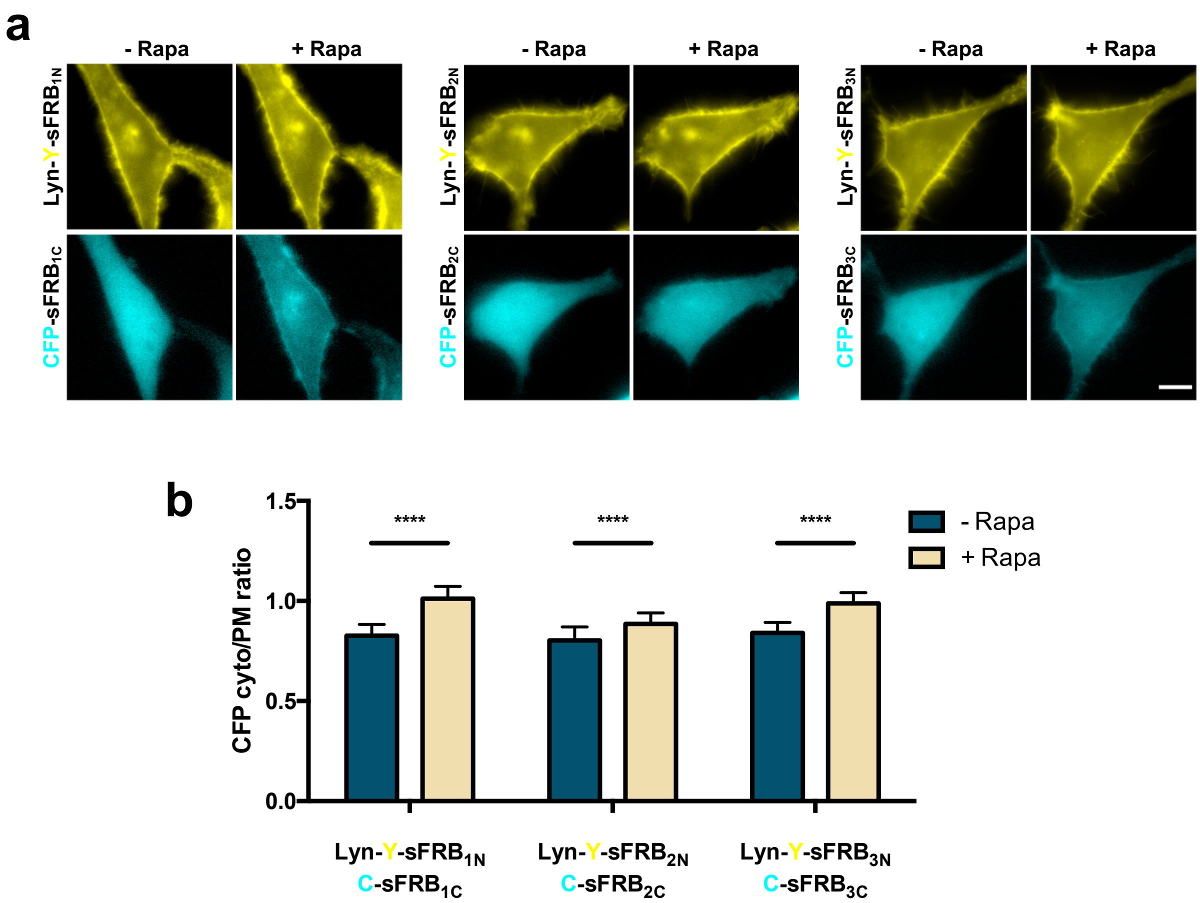

### Supplementary Figure 4

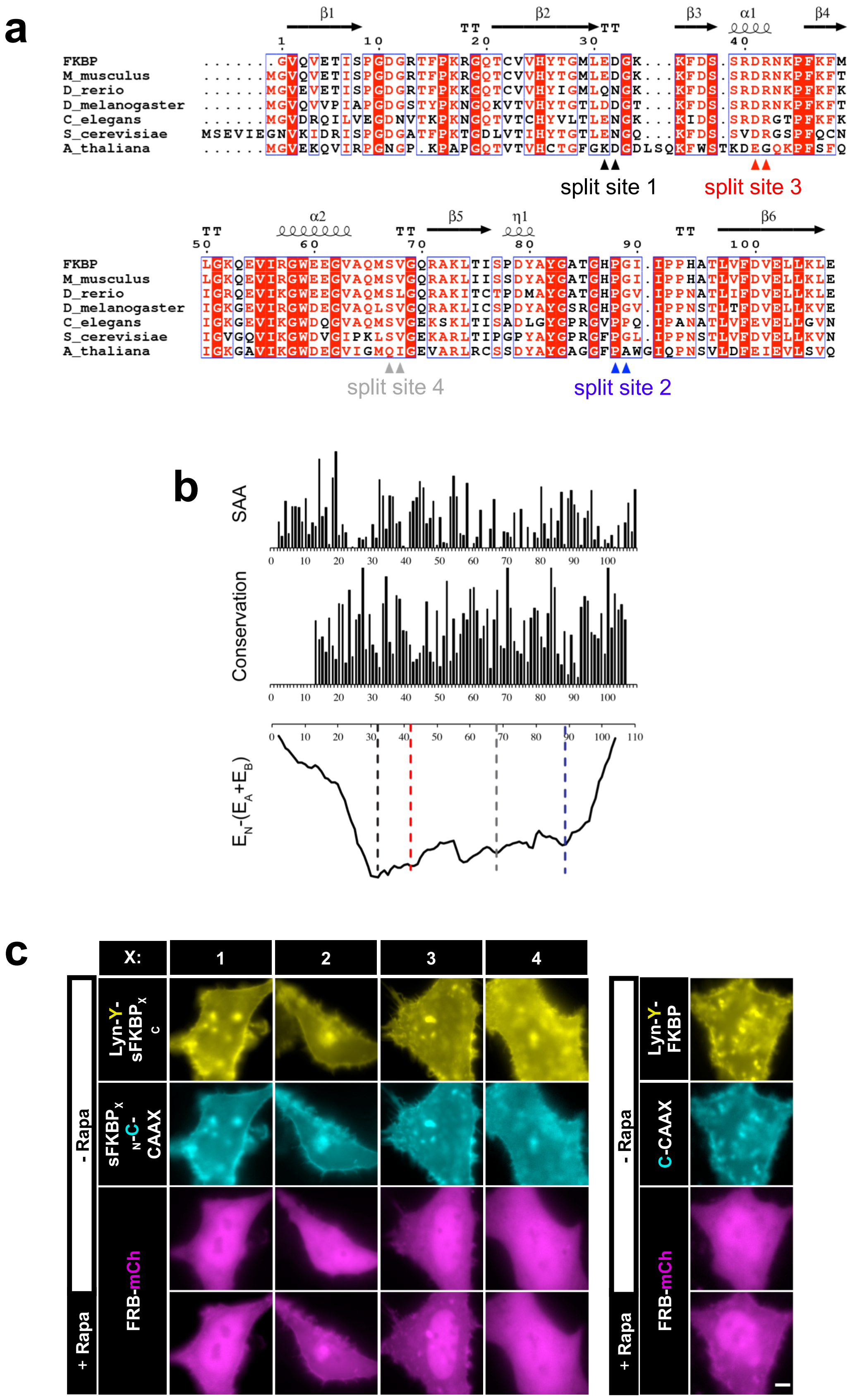

### Supplementary Figure 5

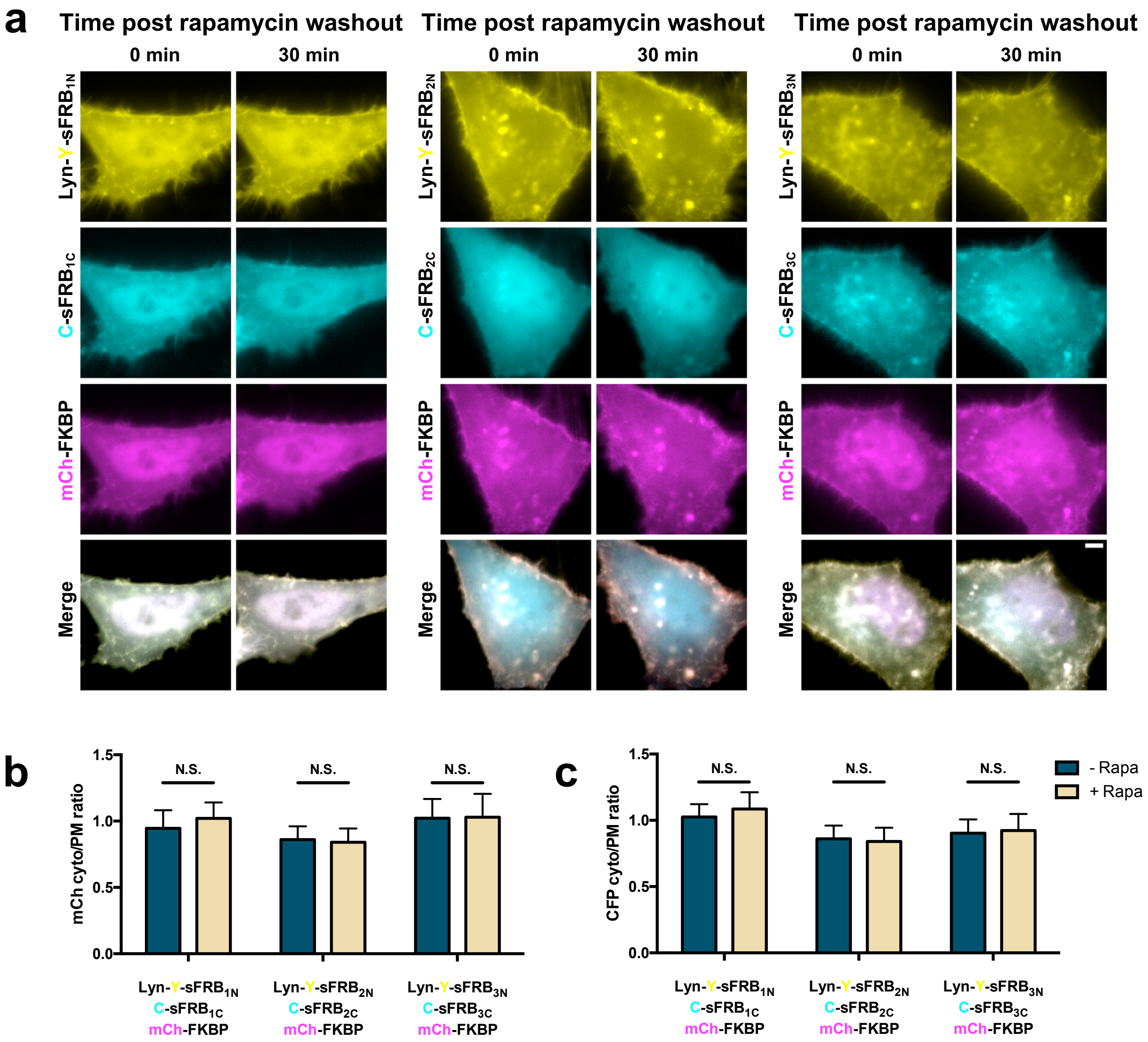

### Supplementary Figure 6

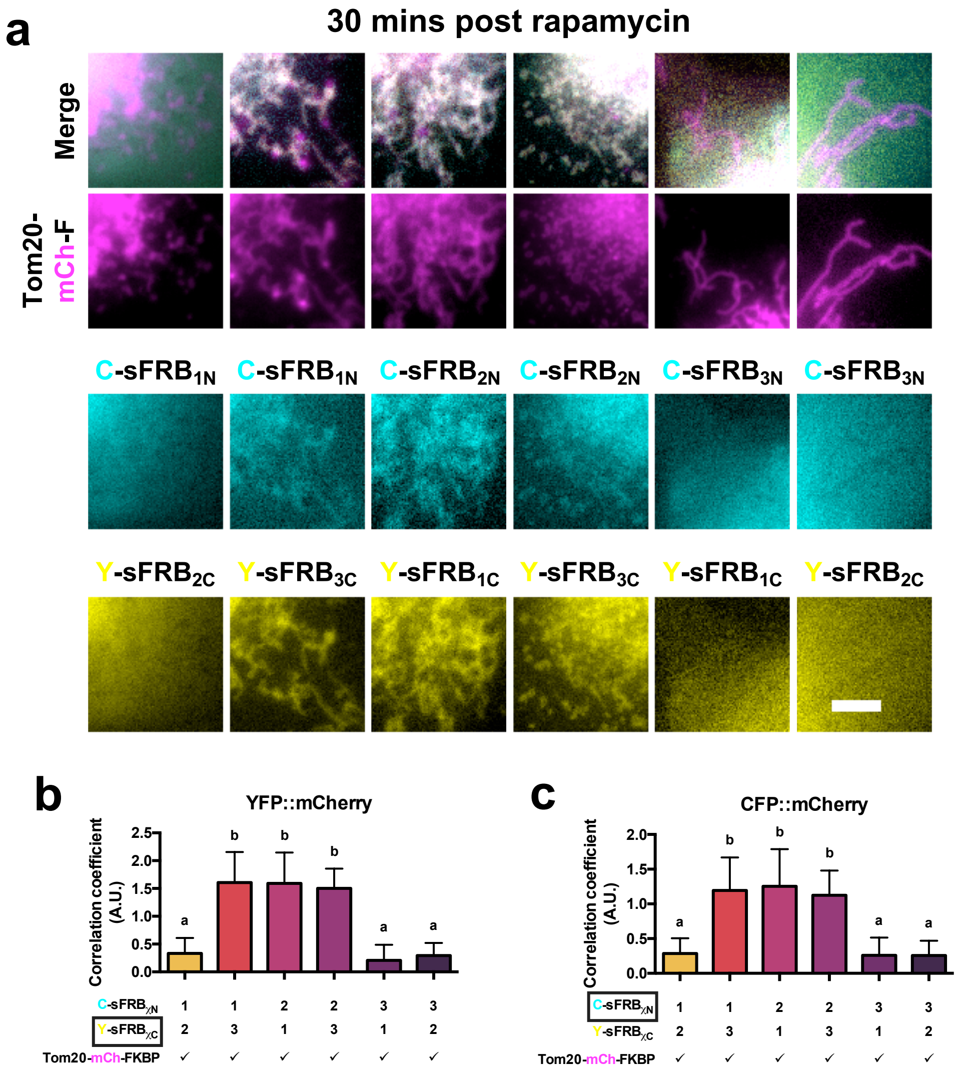

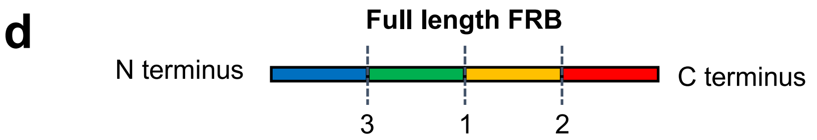

| Part 1 | Part 2 | Illustration | Overlap | Functional? |
|---|---|---|---|---|
| $sFRB_{1N}$ | $sFRB_{2C}$ | | No | No |
| $sFRB_{1N}$ | $sFRB_{3C}$ | | Yes | Yes |
| $sFRB_{2N}$ | $sFRB_{1C}$ | | Yes | Yes |
| $sFRB_{2N}$ | $sFRB_{3C}$ | | Yes | Yes |
| $sFRB_{3N}$ | $sFRB_{1C}$ | | No | No |
| $sFRB_{3N}$ | $sFRB_{2C}$ | | No | No |

### Supplementary Figure 7

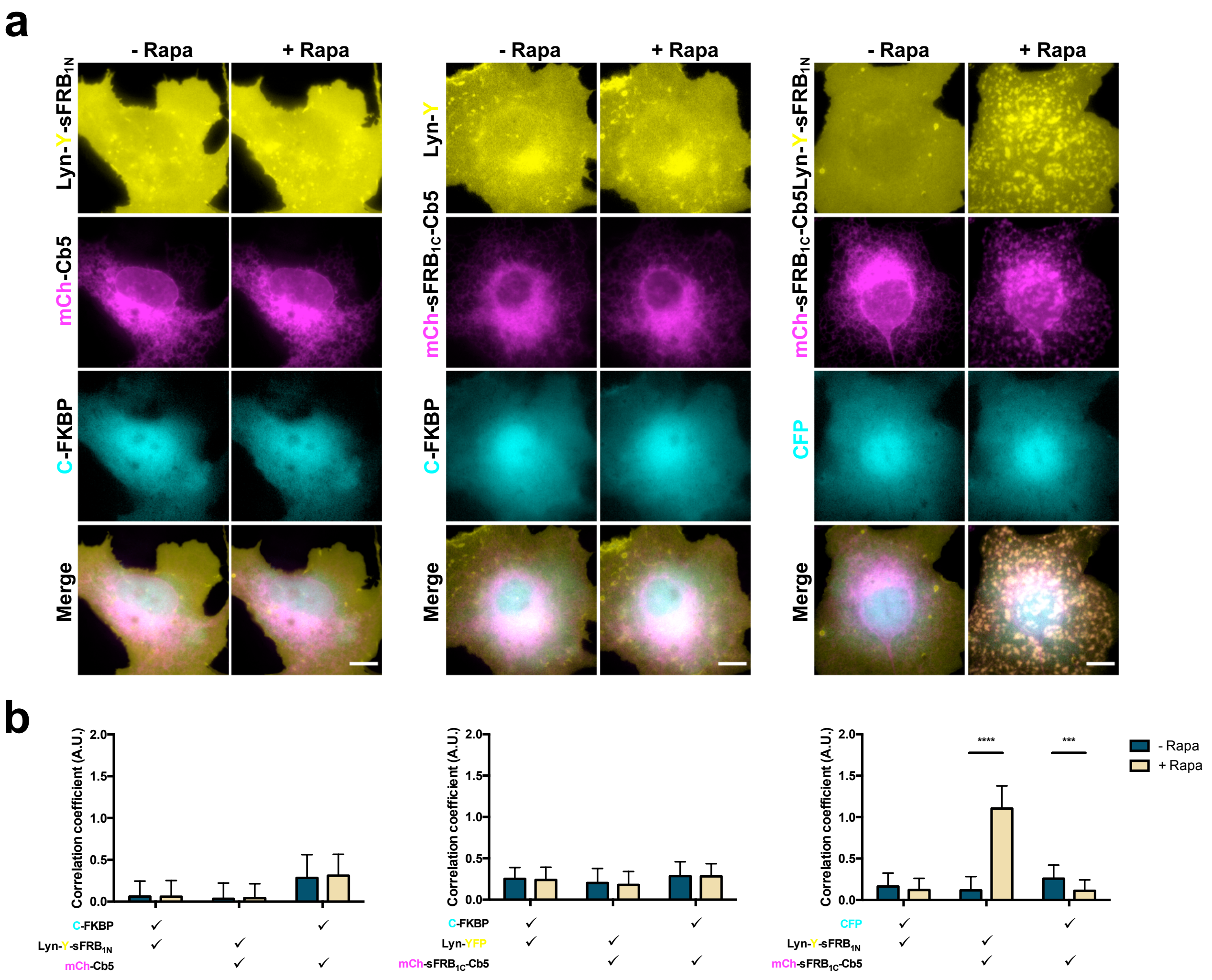

### Supplementary Figure 8

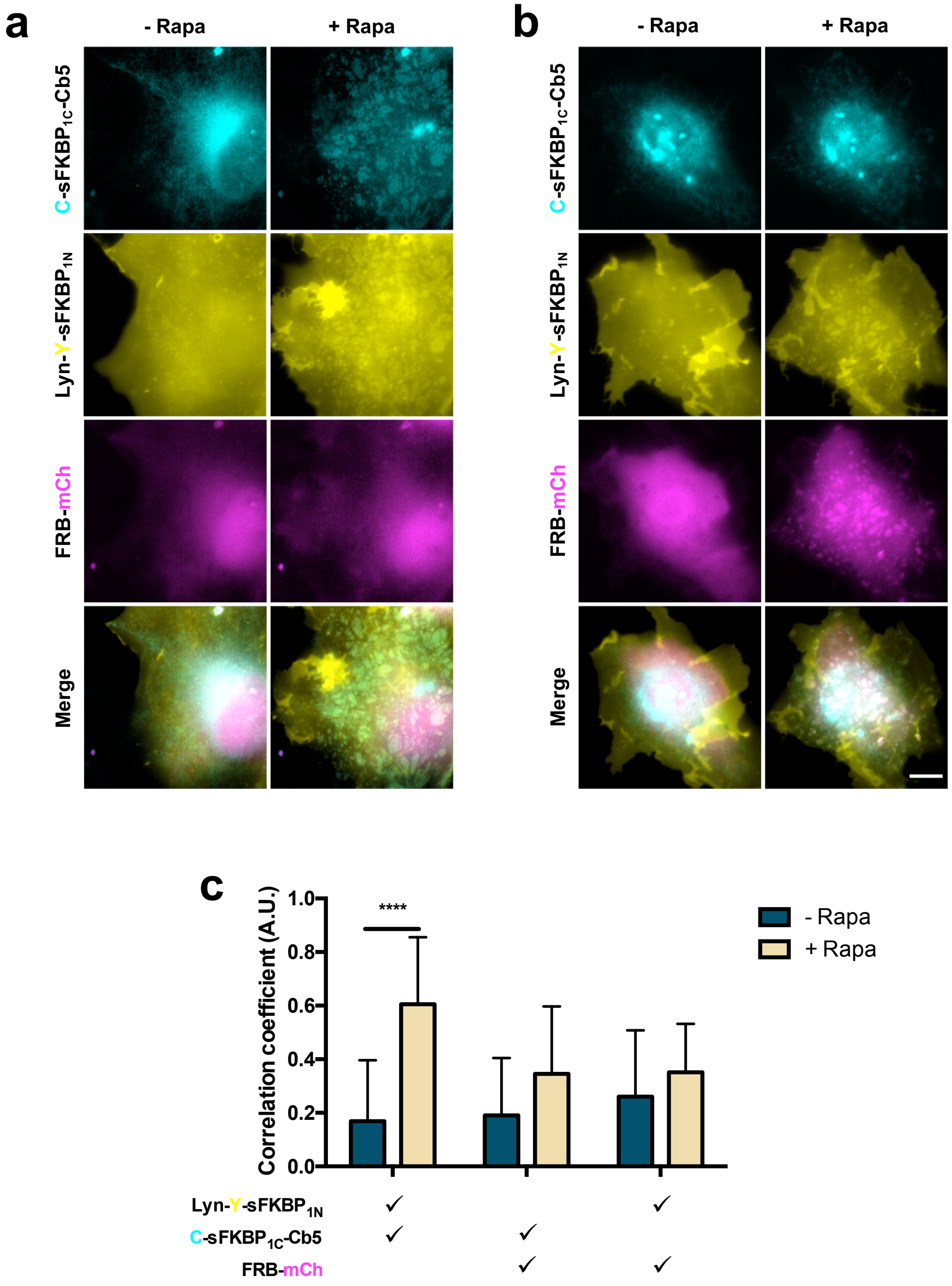

### Supplementary Figure 9

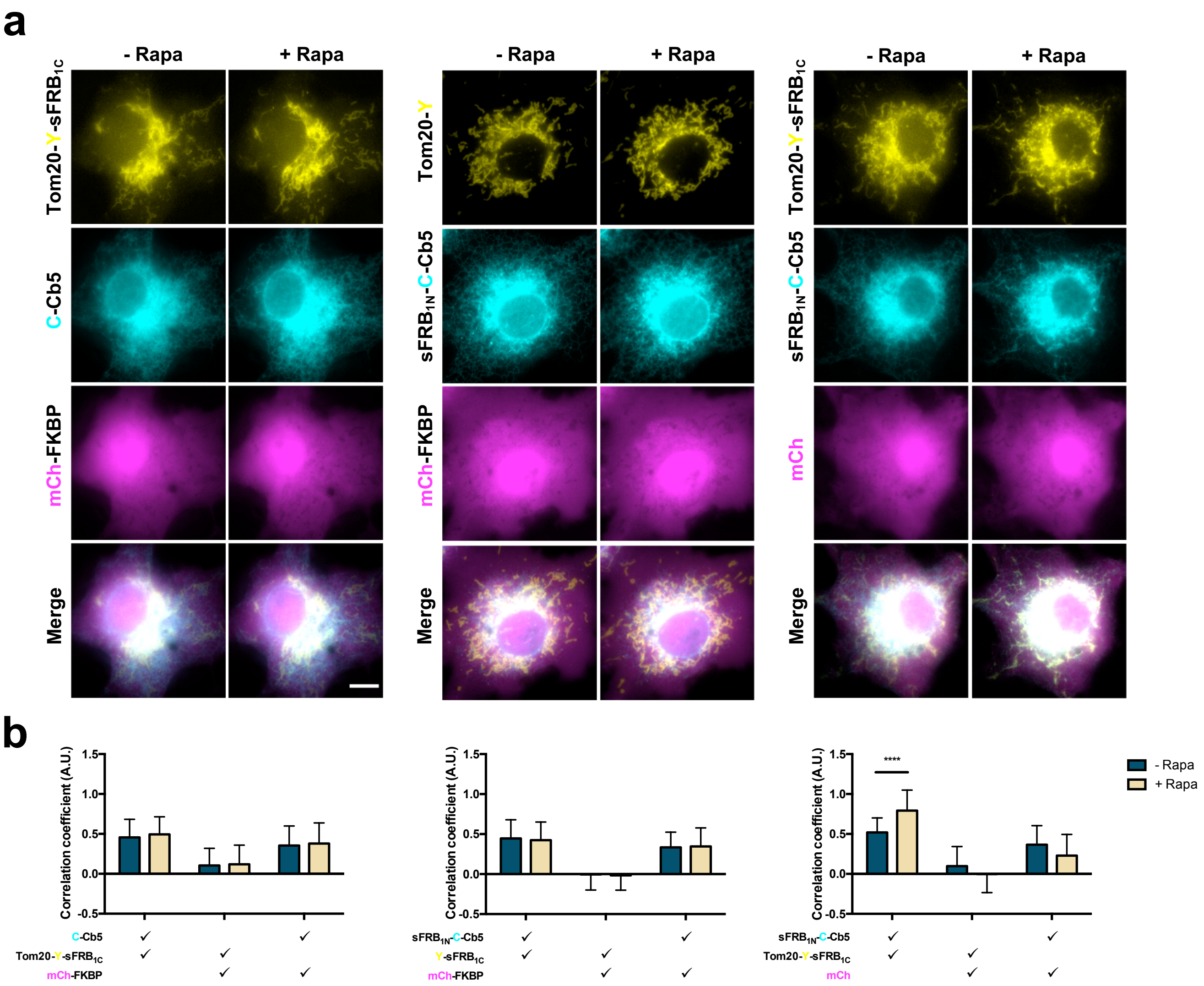

### Supplementary Figure 10

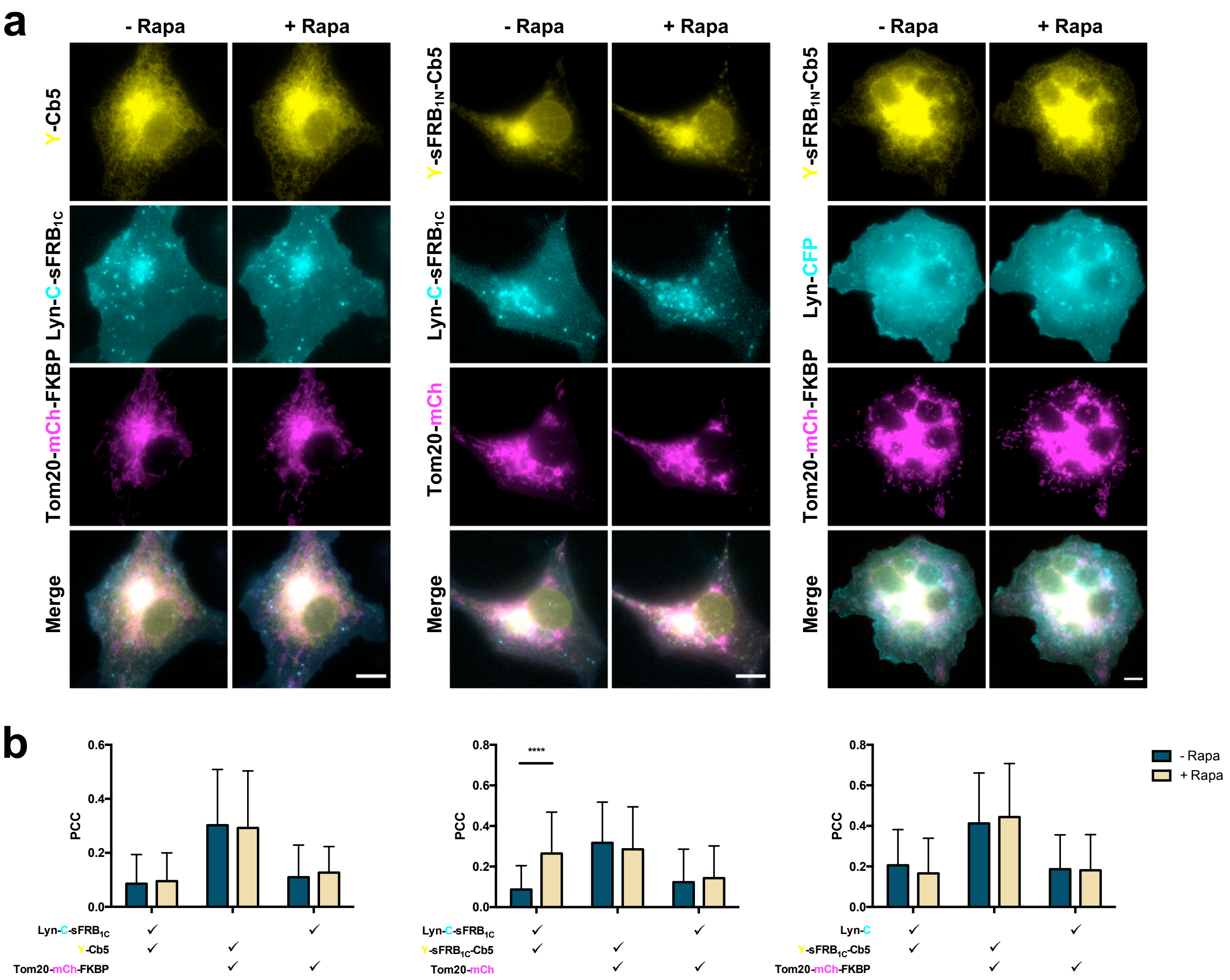

**Supplementary Table 1  Data collection and refinement statistics**

| | FKBP12·rapamycin·FRB (T2098L) | sFKBP$_1$·rapamycin | sFKBP$_1$·rapamycin·FRB (T2098L) |
|---|---|---|---|
| **Data collection** | | | |
| Space group | $P2_1$ | $P2_1$ | $P4_32_12$ |
| Cell dimensions | | | |
| *a*, *b*, *c* (Å) | 57.04, 67.42, 69.79 | 107.03, 48.14, 108.73 | 127.55, 127.55, 278.56 |
| α, β, γ (°) | 90.00, 107.20, 90.00 | 90.00, 106.52, 90.00 | 90.00, 90.00, 90.00 |
| Resolution (Å) | 47.45–2.20 (2.27–2.20) | 48.14–2.92 (3.10–2.92) | 48.65–3.11 (3.23–3.11) |
| $R_{merge}$ | 0.146 (0.846) | 0.109 (0.669) | 0.349 (2.477) |
| $I / \sigma I$ | 9.7 (2.5) | 7.4 (1.8) | 9.5 (1.5) |
| Completeness (%) | 96.6 (100.0) | 100.0 (100.0) | 100.0 (100.0) |
| Redundancy | 6.9 (6.5) | 3.8 (3.8) | 14.8 (15.0) |
| **Refinement** | | | |
| Resolution (Å) | 47.45–2.20 (2.26–2.20) | 48.14–2.92 (3.00–2.92) | 48.65–3.11 (3.19–3.11) |
| No. reflections | 173255 | 89147 | 623820 |
| $R_{work}$ / $R_{free}$ | 0.198 / 0.258 | 0.208 / 0.298 | 0.236 / 0.278 |
| No. atoms | | | |
| Protein | 3259 | 7196 | 13335 |
| Ligand/ion | 157 | 136 | 270 |
| Water | 308 | 154 | 191 |
| *B*-factors | | | |
| Protein | 28.63 | 57.92 | 71.42 |
| Ligand/ion | 25.84 | 43.08 | 54.14 |
| Water | 35.54 | 37.94 | 39.77 |
| R.m.s. deviations | | | |
| Bond lengths (Å) | 0.002 | 0.005 | 0.002 |
| Bond angles (°) | 1.157 | 1.390 | 0.706 |
| **PDB ID** | 6M4U | 6M4V | 6M4W |

Values in parentheses are for the highest-resolution shell. One crystal was used to determine each structure.

**Supplementary Video 1**
Timelapse epifluorescence images of Cos-7 cells showing recruitment of cytosolic FKBP (magenta) to ER-plasma membrane junction sites with CIT upon rapamycin addition. ER in cyan, plasma membrane in yellow, time in mm:ss, scale bar 10 µm.

**Supplementary Video 2**
Timelapse epifluorescence images of Cos-7 cells showing recruitment of cytosolic FRB (magenta) to ER-plasma membrane junction sites with CIT upon rapamycin addition. ER in cyan, plasma membrane in yellow, time in mm:ss, scale bar 10 µm.

**Supplementary Video 3**
Timelapse epifluorescence images of Cos-7 cells showing recruitment of cytosolic FKBP (magenta) to ER-mitochondria junction sites with CIT upon rapamycin addition. ER in cyan, mitochondria in yellow, time in mm:ss, scale bar 10 µm.

**Supplementary Video 4**
Timelapse epifluorescence images of Cos-7 cells showing tri-organellar junction formation between ER, plasma membrane, and mitochondria upon rapamycin addition. Mitochondria in magenta, ER in yellow, PM in cyan, time in mm:ss, scale bar 10 µm.

**Supplementary Video 5**
Timelapse epifluorescence images of Cos-7 cells showing recruitment of INP54P (cyan) to ER-PM junctions resulting in reduced signal intensity of PH-PLCδ (gray). ER in magenta, PM in yellow, time in mm:ss, scale bar 10 µm.

**Supplementary Video 6**
Timelapse epifluorescence images of Cos-7 cells showing recruitment of INP54P D281A (cyan) to ER-PM junctions resulting in no change in signal intensity of PH-PLCδ (gray). ER in magenta, PM in yellow, time in mm:ss, scale bar 10 µm.

**Supplementary Video 7**
Timelapse epifluorescence images of Cos-7 cells showing recruitment of FKBP (cyan) to ER-PM junctions resulting in no change in signal intensity of PH-PLCδ (gray). ER in magenta, PM in yellow, time in mm:ss, scale bar 10 µm.
